# Controlling TCR and CAR activation by targeting LCK recruitment with a first-in-class small-molecule inhibitor

**DOI:** 10.64898/2025.12.10.693420

**Authors:** Nadine M. Woessner, Patricio Chinestrad, Tamina Rückert, Lena Weiß, Marina Zintchenko, Anna-Maria Schaffer, Sara Hartmann, Azin Kiani, Maja Köhn, Maia Cabrera, Frederike A. Hartl, Wolfgang W. Schamel, Natalie Köhler, Pablo Lorenzano Menna, Susana Minguet

**Author notes:** These authors contributed equally (Patricio Chinestrad, Tamina Rückert).

## Abstract

T-cell activation is driven by the recruitment of lymphocyte-specific protein tyrosine kinase (LCK) to the T-cell receptor (TCR), a critical step in initiating immune responses. Existing LCK inhibitors lack specificity because they target the conserved kinase domain shared by Src family kinases, resulting in off-target effects. Here, we introduce a novel strategy to selectively modulate T-cell activation by disrupting the interaction between the SH3 domain of LCK and the receptor kinase (RK) motif of CD3ε. Using computational modeling and high-throughput virtual screening, we identified candidate compounds targeting the SH3(LCK) domain. Functional validation revealed that one compound, C10, selectively disrupted the SH3-RK interaction, leading to reduced TCR-driven activation and proliferation, while sparing activation via alternative receptors and B-cell responses. Moreover, C10 modulated the activity of CD3ε-containing CAR and TRuC T cells, attenuating cytokine production and promoting a central-memory-like phenotype associated with enhanced persistence. These findings establish targeted disruption of LCK recruitment as a viable strategy for fine-tuning T-cell responses and propose SH3(LCK) as a druggable domain with therapeutic potential for autoimmune diseases, graft-versus-host disease, and optimizing CAR T-cell therapies.

## Introduction

T-cell activation is a tightly regulated process initiated by the engagement of the T-cell receptor-CD3 (TCR-CD3) complex, which contains the TCRαβ, CD3εδ, CD3εγ and ζζ dimeric subunits^1^, and triggers intracellular signalling cascades essential for immune responses^2^. A key player in this process is the lymphocyte-specific protein tyrosine kinase (LCK), a Src family kinase (SFK) that phosphorylates immunoreceptor tyrosine-based activation motifs (ITAMs) within the TCR-CD3 complex. These phosphorylation events recruit and activate downstream signalling molecules, ultimately leading to full T-cell activation. However, dysregulated T-cell activation contributes to autoimmune diseases, graft-versus-host disease (GVHD), transplant rejection, and excessive T-cell exhaustion in cancer immunotherapy^3^.

Despite its essential role in TCR signalling, pharmacological modulation of LCK remains a major challenge. All existing inhibitors target the conserved kinase domain of LCK, leading to poor selectivity and off-target effects on other SFKs^4,5^. This lack of specificity limits their clinical utility, particularly for diseases where fine-tuning of T-cell activation is required rather than broad immunosuppression. An alternative approach would be to target a non-catalytic domain of LCK, namely the one which controls LCK’s recruitment to the TCR-CD3 complex.

We recently identified a non-canonical interaction between the SH3 domain of LCK and the receptor kinase (RK) motif in the cytoplasmic tail of CD3ε, which is critical for recruiting LCK to the TCR^6^. Unlike canonical SH3 domain interactions with proline-rich sequences (PRS), the SH3 domain of LCK interacts with CD3ε through a distinct RKxQRxxY motif^6^. This interaction is mediated by the RT and n-SRC loops of the SH3(LCK) domain, which contain residues not conserved in other SFKs, highlighting a unique and specific regulatory mechanism. This previously untapped mechanism thereby offers a novel target for selective T-cell modulation. Unlike kinase inhibitors, targeting this interaction could dampen T-cell activation without completely abolishing signalling. Thus, reducing the risk of systemic immunosuppression. Genetic disruption of the SH3(LCK)-RK interaction has been shown to reduce, but not eliminate, T-cell activation^6^, underscoring the therapeutic potential for targeting this mechanism.

Given the increasing use of engineered T cells in immunotherapy, strategies to fine-tune their activation are of particular interest. We and others have demonstrated the important role of CD3ε in enhancing T cell-based immunotherapies. CD3ε-containing chimeric antigen receptors (CARs) and TCR fusion constructs (TRuCs), which harness the full range of CD3 chains, significantly enhanced anti-tumour efficacy of these therapies^6–12^. However, CAR T cell-therapy can also lead to severe adverse effects, including cytokine release syndrome (CRS), neurotoxicity, and premature exhaustion of the CAR T cells due to excessive signalling^13^. A pharmacological approach to modulate CAR and TRuC signalling could improve both efficacy and safety, offering a promising strategy for optimizing next-generation T-cell therapies.

In this study, we report a first-in-class small-molecule inhibitor, C10, that selectively disrupts the interaction between the SH3 domain of LCK and the RK motif of CD3ε, thereby impairing LCK recruitment to the TCR without affecting its kinase activity. Through computational modeling and high-throughput virtual screening, we identified several SH3(LCK)-targeting candidates, of which C10 uniquely attenuated TCR-induced T-cell activation while sparing alternative activation pathways. C10 also dampened allogeneic T-cell responses, modulated CAR and TRuC T-cell activation, and promoted a central-memory-like CAR T-cell phenotype. By targeting a novel mechanism of LCK recruitment^6^, our findings establish SH3(LCK) as a druggable domain for selective T-cell modulation and offer a promising strategy for immune-mediated diseases and CAR T-cell optimization.

## Materials and Methods

### CD3ε-SH3(LCK) interaction prediction

To predict the key residues in the interaction between CD3ε and SH3(LCK), the AutoDock CrankPep tool was used^14^. It allows to reliably dock of flexible peptides up to 20 amino acids. The docking box dimensions were 24 Å by 20 Å by 24 Å with a grid spacing of 1 Å and was centered around the previously described interaction sites between CD3ε and SH3(LCK)^6^.

### Docking and interaction characterization

In order to identify candidates with high affinity for the SH3(LCK) domain, a High Throughput Virtual Screening (HTVS) was performed. The AutodockVina software^15,16^ was used as the docking software. The docking was centered around the druggable interaction surface in the region of SH3(LCK) domain that interacts with CD3ε. The docking box dimensions were 16 Å by 14 Å by 18 Å with a grid spacing of 1 Å. The Enamine Advanced Collection, a database with over 500.000 drug-like compounds was used in the screening (https://enamine.net/compound-collections/screening-collection/advanced-collection). The preparation of the compounds for docking was performed using the AutoDockTools4 package^17^. For the parallelization and distribution of the docking tasks to our high performance computing cluster, the in-house developed Molecular Sniper software^18^ was used. For the best candidates, the interactions established between them and the pocket of SH3(LCK) were calculated using the Protein-Ligand Interaction Profiler (PLIP) software^19^.

### Fingerprints

For each candidate, E3FP 3D, MACCS Keys^20^, Morgan 2D, and Murcko Scaffold 2D fingerprints were computed. The 4096-bit E3FP 3D fingerprints were generated using the E3FP software^21,22^. The 166-bit Molecular ACCess System (MACCS) Keys and the 1024-bit Morgan 2D fingerprints were calculated using the RDKit toolkit^23^. In addition, a 1024-bit Morgan 2D fingerprint derived from the Murcko scaffold of each candidate was also computed using RDKit. Fingerprint similarity was assessed using the Tanimoto coefficient, as implemented in RDKit, where a value of 1 indicates identical fingerprints.

### Prediction of ligands physicochemical parameters

The physicochemical parameters of each candidate were predicted using the SwissADME web server^24^. The SwissADME web tools suite is specifically designed to predict ADME properties of chemical compounds.

### Sequence and Structural Comparison

Amino acid sequence comparison between human and murine SH3(LCK) was performed using BLASTp (NCBI)^25^. The murine SH3(LCK) structure was obtained from the AlphaFold Protein Structure Database^26–28^. Structural alignment between the murine SH3(LCK) AlphaFold model and the experimentally determined human SH3(LCK) structure was carried out using MAMMOTH^29^.

### CD25 upregulation and IFNγ production

0.2×10^6^ splenocytes were stimulated at 37 °C with 3 μg/ml plate-bound anti-mCD3ε antibody (145-2C11, Biolegend), or with antigen-presenting cells (APCs) derived from CD3ε^-/-^ mouse spleens and loaded with 10 ng/ml OVA, Q4R7 or Q4H7 peptide (for 3-4 h) in a ratio of 1:1 in the presence of 0.33% DMSO or the indicated compounds dissolved in DMSO. After 48 h, cells were stained with anti-mCD4-PE-Cy7 (RM4-5, eBioscience), anti-mCD8-FITC (Invitrogen) and anti-mCD25-APC (PC61.5, eBioscience) antibodies. CD25 upregulation was assessed by flow cytometry. Alternatively, the supernatant of the stimulated cells was collected after 48 h of stimulation and IFNγ secretion was assessed using a commercial ELISA kit (eBioscience).

### T-cell proliferation

0.5×10^6^ splenocytes were stained with 1 μM CellTrace Violet (ThermoFisher) and stimulated at 37 °C with 3 μg/ml plate-bound anti-mCD3ε antibody (145-2C11) or 10 ng/ml PMA and 250 ng/ml ionomycin. After 72 h, the cells were stained with anti-mCD4-PE-Cy7 (eBioscience/Biolegend) and anti-mCD8-FITC/PE-Cy7 (Invitrogen/Biolegend) antibodies. Alternatively, freshly thawed human peripheral blood mononuclear cells (PBMCs) were pre-activated with 2 μg/ml phytohemagglutinin (PHG, Sigma) for 48 h in the presence of 50 ng/ml IL2 (Peprotech) and then cultured for 5-7 days in the presence of 10 ng/ml IL2 before being used for an experiment. Then, 0.5×10^6^ cells were stained with 1 μM CellTrace Violet and stimulated at 37 °C with 3 μg/ml plate-bound anti-hCD3ε antibody (UCHT1) or 100 ng/ml IL2 for 96 h. Alternatively, 0.3×10^6^ Jurkat (JK) cells were stained with 1 μM Cell Trace Violet and incubated in 10% FCS RPMI or 0.1% FCS RPMI (negative control) for 72 h. Human cells were stained with anti-hCD8-APC (Beckmann Coulter) and anti-hCD4-PE-Cy7 (Biolegend) antibodies. Cells were incubated with 0.33% DMSO or the test compounds (C1, C3–C10, or PP2) dissolved in DMSO. T-cell proliferation was determined by the extent of CellTrace Violet dye dilution measured by flow cytometry (Gallios Beckman Coulter/Attune).

### Toxicity

6-well plates were pre-coated with 1 μg/ml anti-hCD3ε (UCHT1) and 1 μg/ml anti-hCD28 antibody for 1 h at 37 °C. Frozen PBMCs from healthy donors were thawed and 1×10^6^ cells/ml were plated into wells in the presence of 100 ng/ml of IL2 (Preprotech). 3 days later, 0.2×10^6^ cells per sample were incubated with 0.33% DMSO or C6, C8, C10, PP2 or A77 dissolved in DMSO for 24 h. The percentage of apoptotic and dead cells was determined with AnnexinV-FITC staining (BD Pharmingen).

### B-cell proliferation

0.3×10^6^ MACS-purified untouched splenic B cells (Miltenyi Biotec) from C57BL/6 mice were stained with 1 μM CellTrace Violet (ThermoFisher) and stimulated at 37 °C in 96-well plate with 10 μg/ml anti-IgM (Fab’2) antibody and 5 ng/ml IL-4 or with 2.5 μg/ml LPS in the presence of 0.33% DMSO or 8 μM of C6, C8, C10 dissolved in DMSO. Cells were stained with anti-CD19-PE antibody (Biolegend) as B cell marker. B-cell activation was assessed by flow cytometry after 24 h of stimulation by staining with anti-CD86-PE (Biolegend) and anti-MHCII-APC (Biolegend) antibodies, while proliferation was determined by the extent of CellTrace Violet dye dilution measured by flow cytometry after 72 h of incubation (Attune).

### Allogeneic T-cell activation

Splenic T cells from C57BL/6 mice were MACS-purified using the Pan T cell isolation kit (Miltenyi Biotec), labeled with 1 μM CellTrace Violet (ThermoFisher) and co-cultured with LPS-stimulated, allogeneic bone marrow-derived dendritic cells (BMDCs) from BALB/c mice at a cell ratio of BMDC:T cells = 1:10 in a 96-well format. The cells were treated with 0.33% DMSO or C6, C8 or C10 dissolved in DMSO and after 72 h cells were stained with anti-mCD69-PE, anti-mCD4-APC and anti-mCD8-PE-Cy7 antibodies (all Biolegend). T-cell proliferation was determined by the extent of CellTrace Violet dye (ThermoFisher) dilution measured by flow cytometry (LSR Fortessa). T cell viability was measured using the Zombie NIR Fixable Viability Kit (Biolegend).

### Intracellular calcium mobilization

For Ca^2+^ influx measurement, 0.5×10^6^ cells per sample were resuspended in medium containing 1% FCS and incubated for 30 min at 37 °C with 5 μg/ml of Indo-1 and 0.5 μg/ml of pluronic F-127 (Molecular Probes). After washing, the cells were kept in the dark on ice. Right before measurement, the cells were pre-warmed 5 min at 37 °C in the presence of 0.08% DMSO or 8 μM C6, C8 or C10 dissolved in DMSO. The Ca^2+^ response was induced by addition 3 μg/ml anti-mCD3ε (145-2C11, Biolegend) or 10 or 0.5 μg/ml anti-hCD3ε (UCHT1) antibody after 60 s of baseline measurement. The change of the ratio of Indo-bound versus Indo-unbound was measured with a MACSQuantX Flow Cytometer (Miltenyi Biotech). Data were normalized to the baseline with FlowJo software version 9.3.2.

### Kinase assay

2 μg hcytCD3ε-GST (GST-fusion protein of the human cytoplasmic tail of CD3ε) or hcytζ-GST (GST-fusion protein of the human cytoplasmic tail of ζ) and 17 μM ATP were incubated in 1x kinase buffer (40 mM HEPES, 10 mM MgCl_2_, 3 mM MnCl_2_) in the presence or absence of 50 ng LCK-GST (ENZO; BML-SE356-005) with 1% DMSO or 100 μM C6, C8, C10 or A77 dissolved in DMSO for 15 min at 30 °C. Reaction was stopped by adding 5x non-reducing sample buffer and boiling the sample for 5 min at 95 °C.

### Cell stimulation for immunoblotting

3×10^6^ Jurkat cells per sample were incubated with 0.33% DMSO or 8 μM C6, C8, C10 or A77 dissolved in DMSO for 1 h at 37 °C. Afterwards, cells were left unstimulated or stimulated with 5 μg/ml anti-hCD3ε (UCHT1) antibody, and then lysed in 80 μl EMBO lysis buffer (20 mM Tris-HCl (pH 8), 137 mM NaCl, 2 mM EDTA, 10% glycerol, protease inhibitor cocktail (Sigma), 1 mM PMSF, 5 mM iodoacetamide, 0.5 mM sodium orthovanadate, 1 mM NaF and 0.3% Brij96V) for 30 min at 4 °C. Samples were stored at −20 °C with 5x non-reducing sample buffer until further use.

### Immunoprecipitation (IP) and pull down (PD)

For the IP, 60×10^6^ Jurkat (JK) cells were treated with 0.33% DMSO or 8 μM of C6, C8, C10 dissolved in DMSO and incubated for 1 h at 37 °C. Cells were either left unstimulated or stimulated with 5 μg/ml anti-CD3ε antibody (UCHT1) for 5 min at 37 °C and then lysed in 1 ml EMBO lysis buffer (see above) for 1 h at 4 °C. 10 μl of anti-pY antibody 4G10-coupled agarose beads (Merck Millipore) were added to 900 μl of the lysate and incubated over night at 4 °C. For the PD, 55×10^6^ JK cells were resuspended in 1 ml of EMBO lysis buffer and lysed for 1 h at 4 °C. 10 μl of cytCD3ε-GST-coupled sepharose beads were incubated with the lysate overnight in the presence of 1% DMSO, or 100 μM of C6, C8 or C10 dissolved in DMSO. In all cases, beads were washed 3x with 500 μl EMBO washing buffer (20 mM Tris-HCl (pH 8), 137 mM NaCl, 2 mM EDTA, 10% glycerol, 0.3% Brij96V) and stored in 20 μl 5x non-reducing sample buffer at −20 °C until further use.

### SDS PAGE and Immunoblotting

Samples were loaded on 12% SDS gels and subjected to SDS-PAGE separation and transferred to PVDF membranes for immunoblotting by semi-dry transfer (18 V; 1 h). Proteins were analyzed by immunoblotting with anti-ζ (self-made), anti-ZAP70 (Cell Signalling), anti-pζ (pY142, Sigma), anti-GST (Bethyl), anti-LCK (Santa Cruz), anti-pLCK (pY505, Cell Signalling), anti-pLCK (pY394(LCK)/pY416(Src), Cell Signalling), anti-GAPDH (Sigma), anti-pCD3ε (pY1, self-made) and anti-NCK1 (Cell Signalling) antibodies. Quantification of the band intensities was performed with the ImageQuantTL software (GE Healthcare) after chemiluminescence detection. Band intensity was set to 1 for the DMSO control.

### *In situ* proximity ligation assay (PLA)

1×10^5^ cells per sample were starved in 0% FCS RPMI and rested on diagnostic microscope slides (Thermo Fisher Scientific) for 1 h at 37 °C. Cells were left unstimulated or stimulated with 5 μg/ml anti-hCD3ε (UCHT1) for 5 min at 37 °C in the presence of 0.33% DMSO or 8 μM of the indicated compounds. Cells were then fixed with 2% PFA for 15 min, permeabilized with 0.5% saponin for 30 min. Afterwards, cells were blocked according to the manufacturer’s instructions and stained with the Duolink kit (Olink Bioscience) with goat anti-CD3ε (Everest Biotechnology) and mouse anti-LCK (3A5, Santa Cruz Biotechnology). Nuclei were stained with DAPI (Roth). A total of 5-7 images (on average 700 cells) per sample were taken with a 60X objective of a confocal microscope (Nikon C2) and analyzed with BlobFinder^30^.

### Generation of lentiviruses

10^7^ HEK293T cells were plated on a 15 cm plate in 20 ml of DMEM and incubated at 37 °C and 7.5% CO_2_. After 24 h, the medium was changed, and the cells were transfected with the respective constructs and the packaging plasmids pMD2.G (envelope) and pCMVdR8.74 (gag/pol) using PEI transfection. The virus-containing supernatant was collected 24 and 48 h after transfection and was concentrated with a 10% sucrose gradient (supplemented with 0.5 mM EDTA) centrifugation for 4 h at 10,000 xg and 6 °C. After centrifugation, supernatant was discarded and the virus pellet was resuspended in 90 μl of 0% FCS RPMI and stored at −80 °C.

### Primary human T-cell activation, lentiviral transduction and expansion

Isolated PBMCs from healthy donors (ethics approval no. 22-1275-S1) were thawed and resuspended in medium supplemented with 50 ng/ml recombinant human IL2 (PeproTech) and activated with 1 μg/ml anti-hCD3ε and anti-hCD28 antibodies. At 48-72 h, the remaining PBMCs were mostly T cells (>99 %). Primary human T cells were then lentivirally transduced using spin infection in the presence of 5 μg/ml protamine sulfate (Sigma) and 50 ng/ml IL2 with a multiplicity of infection (MOI) of 4. Transduced T cells were checked for CAR/TRuC expression 5-7 days after transduction using a biotinylated primary goat anti-mouse F(ab’)2 (Invitrogen) antibody followed by streptavidin-APC (BioLegend) staining. Cells were cultured in medium supplemented with 10 ng/ml IL2 for a maximum of 7 days after transduction before use for the indicated experiments.

### CAR and TRuC T-cell killing assay and IFNγ production

For the Bioluminescence-based cytotoxicity assay, firefly luciferase-expressing tumour cells (Nalm6) were plated at a concentration of 2.5×10^5^ cells/ml in 96-well flat bottom plates in triplicates. Then, 75 μg/ml D-firefly luciferin potassium salt (Biosynth) was added to the tumour cells and bioluminescence (BLI) was measured in the luminometer (Tecan infinity M200 Pro) to establish the BLI baseline. Right after, CAR/TRuC T cells were added at a effector-to-target (E:T) ratio of 1:5 and incubated for 24 h at 37 °C. BLI was measured as relative light units (RLUs). RLU signals from cells treated with 1% Triton X-100 indicate maximal cell death. RLU signals from tumour cells without CAR/TRuC T cells determine spontaneous cell death. Percentage of specific lysis (specific killing) was calculated with the following formula: percentage specific lysis = 100 × (average spontaneous death RLU − test RLU) / (average spontaneous death RLU − average maximal death RLU). IFNγ secretion was assessed after 24 h of co-culture by ELISA using the supernatants from co-incubation of Nalm6 and CAR/TRuC T cells at E:T ration of 1:5.

### Resting protocol for CAR T-cell expansion

Two days after transduction, 0.2×10^6^ cells were seeded and incubated with 2, 4 or 8 μM of C10 or a corresponding 0.11% DMSO control for a total of 13 days. Fold induction of living cells was calculated between start of the treatment and end of the treatment. Cells were washed and either left unstimulated or stimulated with Nalm6 target cells for 24 h in the absence of any compound. Afterwards, cells were stained with an anti-CD62L-PECy7 (Biolegend) and an anti-CD45RA-V450 (Bioscience) antibody to assess CAR T-cell differentation by flow cytometry. In parallel, cells were stained for CD107a (Biolegend) to assess degranulation upon 3 h of target cell encounter by flow cytometry. A luciferase-based killing assay (see above) was used to determine cytotoxic activity over time. In addition, cells were stained with anti-CD69 (Life Technologies), anti-CD25 (Invitrogen) and anti-4-1BB (eBioscience) antibodies to assess upregulation of the activation markers by flow cytometry after 24 h of stimulation with Nalm6. In addition, the 24 h co-culture supernatants were collected and IL2, IFNγ and TNF secretion were assessed using commercial ELISA kits (eBioscience).

### Luciferase-based proliferation assay of leukemic B-cell lines

0.3×10^6^ Luciferase-expressing B cells were seeded in the presence of 37.5 μg/ml firefly D-luciferin. Nalm6 expressing the oncogenic BCR IGLV3-21R110^31^ were included in the study to test the effect of the compounds on oncogenic BCR signalling. Cells were either left untreated or treated with 0.33% DMSO or 8 μM of C6, C8 or C10. Bioluminescence (BLI) was measured in the luminometer (Tecan infinity M200 Pro) after 24, 48 or 72 h.

### Quantification and statistical analysis

Statistical parameters including the exact number of samples (n) and precision measures (means and ± SD) as well as statistical significance are reported in figures and figure legends. Data sets were assessed for normal distribution with the Shapiro-Wilk normality test, which is recommended for biological samples with a sample size <50 ^32^. Data sets which passed the Shapiro-Wilk test were analysed with the ordinary one-way ANOVA or the two-way ANOVA with Dunnett’s multiple comparison test as indicated (two-tailed). Data were judged to be statistically significant when P < 0.05 and the exact P value is indicated in the graphs. Statistical analysis was performed in GraphPad Prism software version 10.1.1.

### Data availability

The authors declare that all relevant data supporting the findings of this study are available within the paper and its supplementary information files.

## Results

### Docking-based virtual screening identifies novel small-molecule inhibitors targeting the SH3(LCK) domain

The unique and non-canonical interaction between the receptor kinase (RK) motif of CD3ε and the SH3 domain of LCK plays an important role in TCR activation^6^. We hypothesized that disrupting this protein-protein interaction (Fig. 1a) could serve as a precise strategy to modulate TCR signalling. To explore this, we first generated a computational model of the human SH3(LCK) domain bound to the cytoplasmic tail of CD3ε (Fig. 1b). This analysis confirmed the key residues that mediate the SH3(LCK)–RK interactions (Fig. 1c): H70, S71, S75, H76, D79, E96, W97, F113, and N114^6^. In line with our objective of disrupting this interaction, we selected this interface region as the target for a docking-based virtual screening of over 500,000 drug-like compounds from the Enamine Advanced Collection. Compounds were ranked based on their predicted binding affinity for SH3(LCK) using the Protein-Ligand Interaction Profiler (PLIP) software^19^, leading to the identification of 10 top candidates (Fig. 1d) predicted to disrupt SH3(LCK)-RK binding. To assess scaffold diversity, we performed chemical similarity analysis using four orthogonal fingerprinting methods (E3FP 3D, MACCS keys, Morgan 2D, and Murcko scaffold 2D). These analyses confirmed that the selected compounds represent distinct chemical families and are not derivatives of a shared scaffold (Supplementary Fig. 1a). We further investigated the drug-like properties of these candidates by evaluating their absorption, distribution, metabolism, and excretion (ADME) profiles using SwissADME *in silico* analysis^24^. The consensus LogP_o/w_ (mean of five calculated partition coefficients between *n*-octanol and water) as a measure of the lipophilicity, demonstrated that all compounds are lipophilic, indicated by the positive value of the ratio (Fig. 1d). In line, the water solubility was assessed as moderate or poor. The compounds, except for C3 and C10, were not predicted to passively permeate the blood-brain barrier (BBB). The drug-likeliness was assessed with the Lipinski filter^33^, showing no violations against its rule-of-five, and the pattern recognition method PAINS (pan assay interference compounds)^34^ predicted no alerts for potentially problematic substructures (Fig. 1d).

**Fig. 1:**
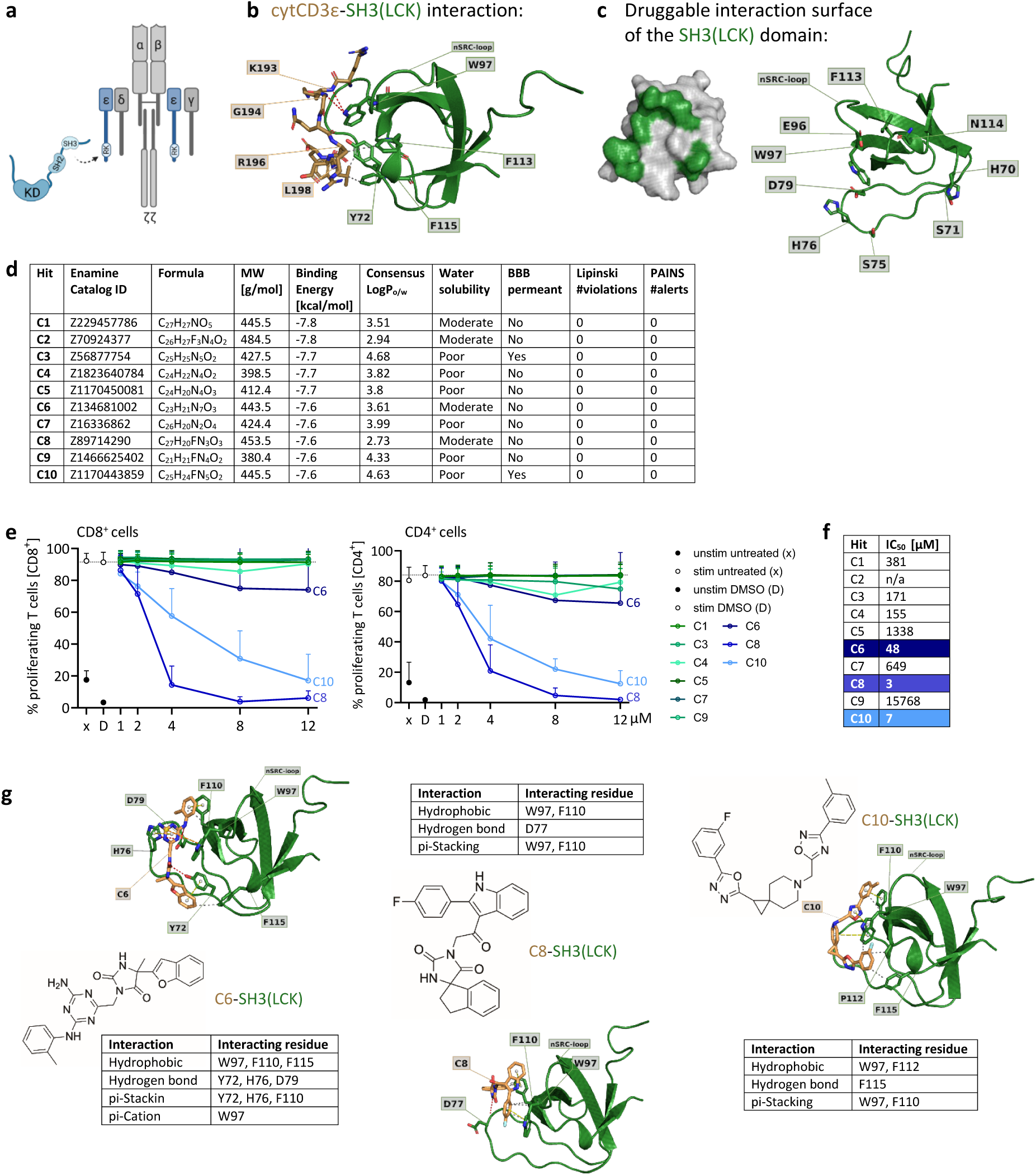
Computational and functional screening identifies small molecules inhibiting T-cell activation. **a**, Schematic representation of LCK recruitment via its SH3 domain to the RK motif of CD3ε, contributing to TCR signalling. **b**, *In silico* model of CD3ε interaction with the SH3(LCK) domain. **c**, Computational modelling identified a druggable interaction surface within the SH3 domain of LCK, which was used for docking simulations with different small molecule compounds. **d**, Top 10 small molecules predicted to bind the druggable pocket of SH3(LCK), including their Enamine Advanced Collection catalogue ID, chemical formula, molecular weight (MW), predicted binding energy to SH3(LCK), and *in silico* ADME parameters: lipophilicity (Consensus Log P_o/w_), water solubility, blood-brain barrier (BBB) permeability, number of Lipinski violations, and PAINS alerts. **e**, Proliferation of CD8^+^ and CD4^+^ primary human T cells was assessed by flow cytometry after stimulation with 5 μg/ml anti-hCD3ε in the presence of 0.33% DMSO or compounds for 96 h. n = 4-6 healthy donors (HD). Means + SD are shown. **f**, IC_50_ values for all compounds calculated based on data from (**e**). **g**, Chemical structures of compounds C6, C8, and C10 (brown) and their predicted interactions with the SH3(LCK) domain (green).

To functionally validate the newly identified compounds, we tested their effects on TCR-induced proliferation of primary human CD4^+^ and CD8^+^ T-cells. The cells were either left untreated (x), treated with 0.33% DMSO (D) or exposed to the individual compounds, and stimulated with an anti-hCD3ε antibody. After 96 h, T-cell proliferation was assessed via flow cytometry. Inhibitory effects were observed for C6, C8, and C10 in both CD4^+^ and CD8^+^ T cells, while the other six tested compounds showed no effect (Fig. 1e). C2 was insoluble in DMSO and was therefore excluded. C6 showed an IC_50_ of 48 μM, while C8 and C10 demonstrated a more potent inhibition with IC_50_ values of 3 and 7 μM, respectively (Fig. 1f). Verifying the results in a proliferation assay using the αβ T cell line Jurkat (JK) confirmed an IC_50_ of C8 and C10 in the low μM range (3 and 9 μM), while C6 did not inhibit proliferation in JK cells (Supplementary Fig. 1b). Therefore, C6 will pose as negative control compound in the following experiments.

A closer analysis of the predicted interactions of the three compounds C6, C8, and C10 with the SH3(LCK) domain revealed that the interactions are mostly mediated by hydrophobic interactions, hydrogen bonds, non-covalent pi-stackings and electrostatic pi-cation interactions (Fig. 1g).

To determine cellular toxicity, we compared C6, C8, and C10 to the commercially available SFK inhibitor PP2 and the LCK inhibitor A-770041 (A77), which both target the kinase domain^35,36^. To this end, we left primary human T cells either untreated (x), treated with DMSO (D), the novel compounds, PP2, or A77 for 24 h and then performed an Annexin V staining to detect apoptotic and dead cells. Treatment of the cells with DMSO as well as treatment with up to 8 μM of C10 or PP2 did not increase the percentage of dead and apoptotic cells compared to the untreated control. Treatment with 8 μM of C8 increased the percentage of dead and apoptotic cells by 10%, and treatment with C6 by 15%, compared to the untreated control (Supplementary Fig. 1c). Thus, we defined these concentrations as suitable to be used in living cells. A77 was highly toxic, inducing 80% cell death at just 2 μM, indicating that it is unsuitable for prolonged T-cell treatment.

Next, we assessed the stability of C6, C8, and C10 at 37 °C by incubating the compounds in culture medium for 3 and 7 days. After incubation, their effects on primary human T-cell proliferation were compared to freshly prepared aliquots. While prolonged exposure to 37 °C reduced their inhibitory activity, C8 and C10 retained 10-20% inhibitory potency even after 7 days, comparable to PP2, which similarly reduced T-cell proliferation by ∼20% after having been kept at 37 °C for 7 days (Supplementary Fig. 1d). These results indicate that C8 and C10 have a stability profile similar to the SFK inhibitor PP2, making them suitable for biological applications.

Overall, through computational modeling and docking-based virtual screening, we identified two novel inhibitors (C8, C10) capable of reducing TCR-mediated T-cell proliferation at a low micromolar range and demonstrated a toxicity and stability profile similar to the well-established SFK inhibitor PP2.

### C8 and C10 suppress TCR activation and allogeneic T-cell responses

To further evaluate the ability of C6, C8, and C10 to modulate TCR-induced activation, we performed a comprehensive set of assays using primary murine splenic T cells. The SH3 domain of murine LCK is identical in amino acid sequence to its human counterpart (Supplementary Fig. 2a), supporting conservation of the binding interface. Structural alignment between the AlphaFold-predicted murine SH3(LCK) and the experimentally resolved human structure revealed a low Root Mean Square Deviation (RMSD) of 0.235 Å, confirming that both domains are structurally indistinguishable (Supplementary Fig. 2b). This supports the use of murine cells for functional testing.

First, we stimulated splenic T cells with an anti-mCD3ε antibody in the presence of DMSO (control), C6, C8, or C10, and T-cell proliferation was measured after 72 hours by flow cytometry (Fig. 2a). The results were consistent with those obtained in human T cells, showing a mild inhibitory effect for C6, a strong effect for C8, and a moderate effect for C10 (Fig. 2a). In a parallel setup, T cells were stimulated with anti-mCD3ε for 48 hours to evaluate the upregulation of the activation marker CD25 by flow cytometry and the production of IFNγ in cell-free supernatants by ELISA (Fig. 2b,c). As expected, C6 had only a mild inhibitory effect, reducing the percentage of CD25^+^ cells by 20% and IFNγ secretion by 10% compared to the DMSO control. In contrast, C8 showed a pronounced, dose-dependent effect, reducing CD25⁺ cells by 40% and IFNγ secretion by 75% at 8 μM. C10 also acted in a dose-dependent manner, decreasing CD25^+^ cells by 50% compared to the DMSO control. Notably, C10 significantly reduced IFNγ secretion (70%) even at 2 μM, indicating a potent effect on cytokine release (Fig. 2b,c). Next, we investigated intracellular calcium mobilization as a more upstream read-out for TCR signalling using Indo-1 staining and flow cytometry analysis. After 60 sec of baseline measurement, splenic T cells were stimulated with an anti-mCD3ε antibody in the presence of DMSO or 8 μM of C6, C8 or C10. Incubation with C6, C8 or C10 resulted in a significant reduction of the percentage of Ca^2+^ responding cells upon TCR stimulation by 25% (C6), 40% (C8), and 50% (C10), compared to the DMSO control (Fig. 2d). This suggests that C8 and C10 interfere with proximal TCR signalling, potentially through inhibition of LCK recruitment.

**Fig. 2:**
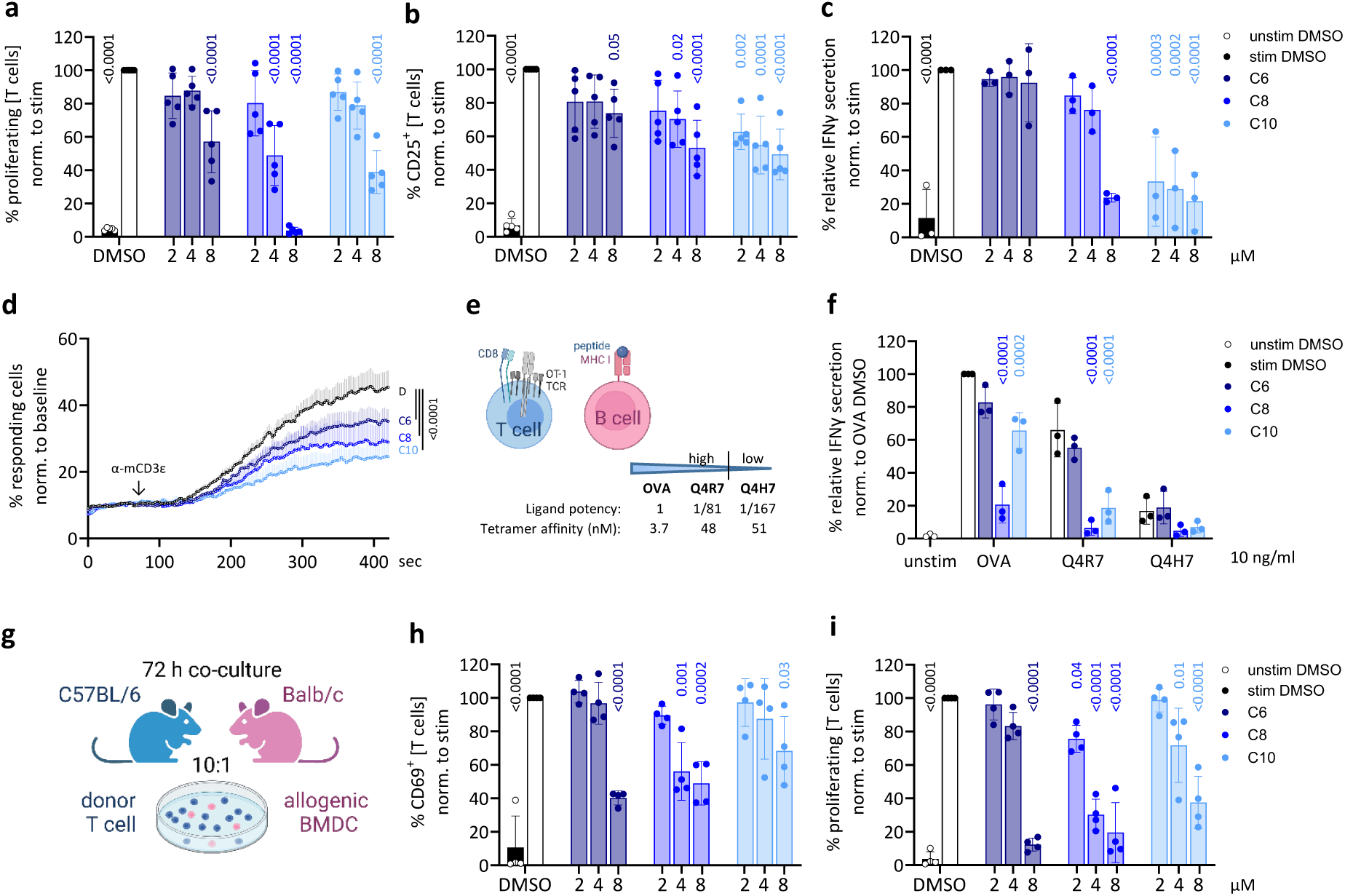
C8 and C10 inhibit TCR-dependent activation and proliferation of primary murine T cells. **a-c**, Splenic T cells were either left unstimulated or stimulated with 3 μg/ml anti-mCD3ε in the presence of 0.33% DMSO, C6, C8 or C10. **a**, T-cell proliferation was evaluated by flow cytometry after 72 h. **b,** The percentage of CD25^+^ T cells was analysed by flow cytometry and **c**, IFNγ production was assessed by ELISA after 48 h. n = 3-5 mice. Means ± SEM are shown. **d**, Calcium influx in splenic T cells upon 3 μg/ml anti-mCD3ε stimulation, measured after 60 s of baseline recording. Cells were treated with 0.33% DMSO or 8 μM of C6, C8, or C10. n = 4 mice. Means + SEM are shown. **e**, OT-1 splenocytes were co-cultured with CD3ε^-/-^ APCs loaded with 10 ng/ml of different OVA peptide derivatives: OVAp (SIINFEKL; high-affinity), Q4R7 (SIIQFERL; mid-affinity), and Q4H7 (SIIQFEHL; low-affinity). **f**, IFNγ production after 48 h of co-culture with 0.33% DMSO or 8 μM of C6, C8, or C10 was assessed by ELISA. n = 3 mice. Data are normalized to OVA-stimulated DMSO control. Two-way ANOVA was used for statistical analysis. **g**, Allogeneic mixed lymphocyte reaction: Splenic donor T cells (C57BL/6) were co-cultured with allogeneic BALB/c bone BMDC at a 10:1 ratio in the presence of 0.33% DMSO, C6, C8, or C10. **h**, The percentage of CD69^+^ cells and **i**, T-cell proliferation were analysed by flow cytometry after 72 h. n = 4 mice. Data are normalized to the stimulated DMSO control. Means ± SD are shown (except d). One-way ANOVA was used for statistical analysis (except f).

We next examined whether these inhibitory effects can be maintained in a more physiological and antigen-specific setting, where the co-receptor CD8 is also recruited due to simultaneous pMHC binding to the TCR and CD8^37^. This would bring CD8-associated LCK to the TCR^38^, potentially reducing the dependency of LCK being recruited directly to the TCR via the SH3(LCK)-RK motif mechanism. To this end, we used the OT-1 TCR transgenic system. OT-1 splenocytes were stimulated with antigen-presenting cells (APCs), derived from spleens of CD3ε^-/-^ mice, and loaded with OVA peptides or derivates exhibiting different affinities for the OT-1 TCR. OVA (SIINFEKL) is a high-affinity ligand, Q4R7 (SIIQFERL) is a mid-affinity ligand, while Q4H7 (SIIQFEHL) is a low-affinity ligand^39^ (Fig. 2e). After 48 h of co-culture, secreted IFNγ was detected by ELISA. All compounds caused a dose-dependent reduction of IFNγ production when T cells were stimulated with any of the three different affinity peptides. C6 reduced IFNγ secretion by 20% (OVA, Q4R7), C8 by 80-90% across all ligands, and C10 by 30% (OVA) and 70-80% (Q4R7 and Q4H7), compared to the DMSO control. These findings indicate that C8 and C10 inhibit TCR activation even in the presence of co-receptor-associated LCK and across a range of pMHC affinities, suggesting their potential to fine-tune T-cell activation in physiological settings.

We next evaluated whether these compounds could suppress allogeneic T-cell activation, a key driver of graft-versus-host disease (GVHD) following allogeneic hematopoietic cell transplantation. Therefore, donor T cells were co-cultured with allogeneic bone marrow-derived dendritic cells (BMDC) in the presence of DMSO, C6, C8 or C10 for 72 h (Fig. 2g). Upregulation of the activation marker CD69 and proliferation was then assessed by flow cytometry. Incubation of the cells with 8 μM of the compounds reduced the percentage of CD69^+^ T cells by 40-60%, compared to the DMSO control (Fig. 2h). The inhibitory effect on T-cell proliferation was even stronger, with a reduction by 60-90% with 8 μM of the compounds (Fig. 2i), demonstrating potent suppression of alloreactive T-cell expansion.

Taken together, these results demonstrate that C8 and C10 suppress TCR-driven activation and T-cell responses – even in co-receptor-dependent and allogeneic settings. These findings highlight their potential as potential immunomodulators for autoimmune diseases and GVHD therapy.

### C10 spares B-cell and IL2-induced activation, supporting TCR specificity

To determine whether the inhibitory effects of our compounds are selective for T cells, we evaluated their impact on B-cell activation. Murine splenic B cells were stimulated either via the B cell receptor (BCR) with an anti-IgM(Fab’2) antibody or via the Toll-like receptor 4 (TLR4) using lipopolysaccharide (LPS) in the presence of 8 μM of C6, C8, or C10. After 24 h, we assessed the upregulation of the activation markers CD86 and MHC-II, and after 72 h, we measured B-cell proliferation via flow cytometry (Fig. 3a,b). Upon BCR stimulation with anti-IgM, C8 reduced the MFI of MHC-II by 20% and both C6 and C8 reduced B-cell proliferation by 40% compared to the DMSO control. Upon TLR4 stimulation with LPS, C6 reduced proliferation by 80%, while C8 reduced it by 35% compared to the DMSO control, suggesting partial off-target effects of C6 and C8 (Fig. 3a,b). Importantly, C10 had no detectable impact on B-cell activation or proliferation, indicating a high specificity for T cells.

**Fig. 3:**
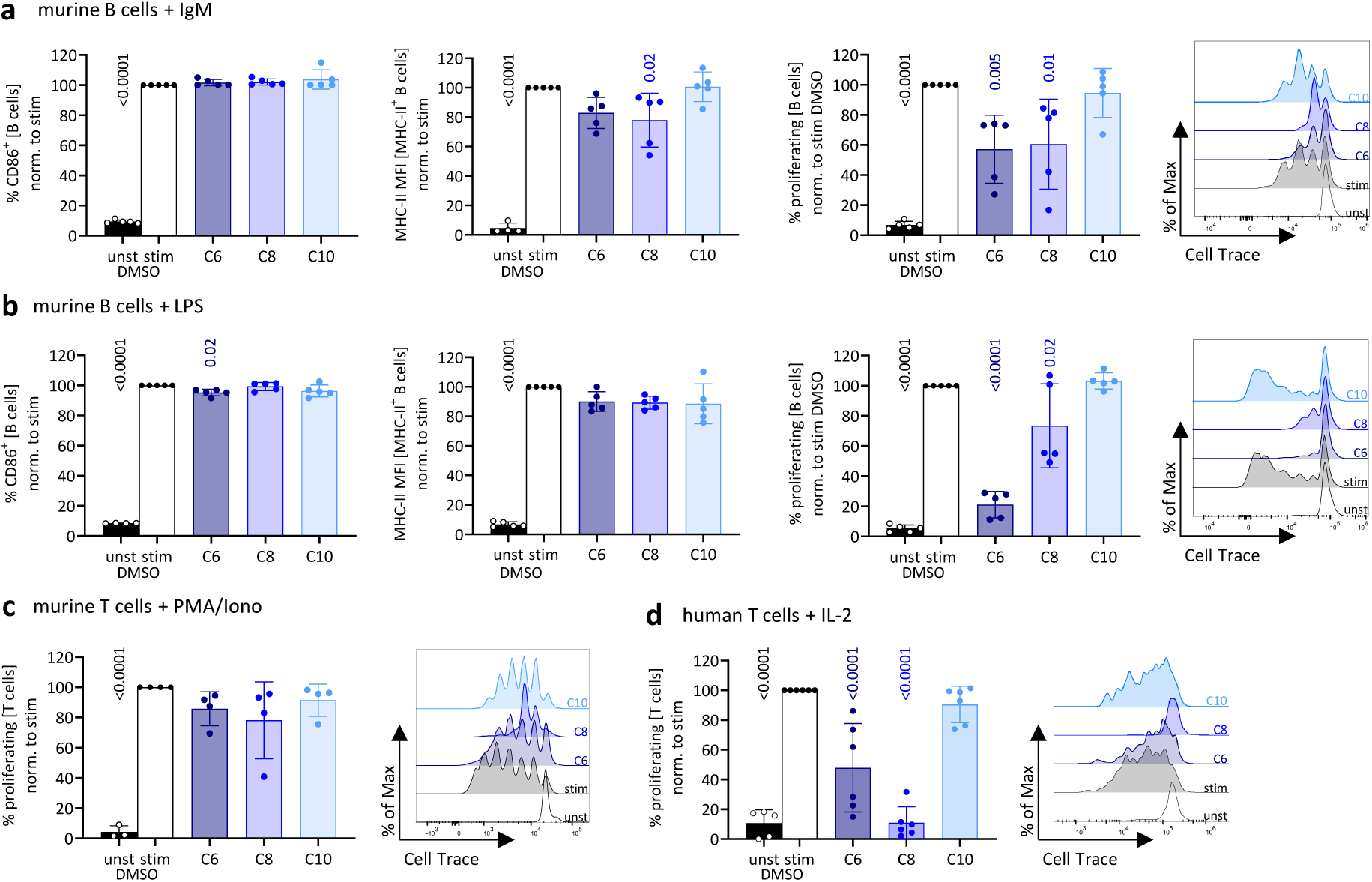
C10 selectively spares B-cell and TCR-independent T-cell activation, confirming both cell type and pathway specificity. Cells were incubated with 0.33% DMSO or 8 μM C6, C8, or C10. **a**, Splenic B were either left unstimulated or stimulated with 10 μg/ml anti-IgM(Fab’2) antibody and 5 ng/ml IL-4 or **b**, 2.5 μg/ml LPS. The percentage of CD86^+^ B cells and the mean fluorescence intensity (MFI) of MHC-II in MHC-II^+^ B cells were assessed after 24 h, while proliferation was evaluated after 72 h by flow cytometry. n = 5 mice. **c**, Splenic T cells were left unstimulated or stimulated with 10 ng/ml PMA and 250 ng/ml Ionomycin; proliferation was assessed after 72 h by flow cytometry. n = 4 mice. **d**, Primary human T cells were left unstimulated or stimulated with 100 ng/ml IL2 for 96 h, and proliferation was assessed by flow cytometry. n = 6 HDs. Means ± SD are shown. One-way ANOVA was used for statistical analysis.

To not only assess cell type specificity, but to also evaluate pathway specificity, we test whether these compounds specifically target TCR-mediated signalling in T cells. We stimulated murine splenic T cells with phorbol-12-myristate 13-acetate (PMA) and ionomycin (Iono). This stimulation bypasses TCR proximal signalling events by directly activating protein kinase C (PKC) and calcium influx. C10 had no significant effect on PMA/Iono-induced T-cell proliferation (Fig. 3c), supporting its specificity for TCR-dependent signalling. C8 exhibited a mild inhibitory effect seen in the representative histogram, suggesting that it may partially interfere with non-TCR-specific downstream activation pathways. Taking into account that PMA/Iono induces strong T-cell activation and might shield possible effects of our compounds, we also investigated IL2-dependent proliferation of primary human T cells. Signalling by the IL2 receptor does not involve LCK, but relies on Janus kinases JAK^40–42^. C10 did not reduce IL2-dependent T-cell proliferation compared to the DMSO control, reinforcing its specificity for TCR signalling and LCK. C8 strongly reduced the percentage of proliferating cells by 80%, indicating that it does not exclusively target the TCR pathway. C6 moderately reduced IL2-dependent T-cell proliferation by 50% (Fig. 3d), confirming that this compound is neither TCR nor T-cell specific.

To exclude potential confounding effects from stereoisomeric variation, we tested the individual enantiomers of C8 and C10. For C8, one enantiomer (C8.a) strongly inhibited TCR-mediated T-cell proliferation (IC₅₀ = 1 μM), while the other (C8.b) was markedly less potent (IC₅₀ = 17 μM) (Supplementary Fig. 3a). C8 enantiomers differentially affected IL2-driven proliferation: C8.a completely abrogated it, whereas C8.b caused a 20% reduction compared to the DMSO control (Supplementary Fig. 3b). In contrast, both enantiomers of C10 inhibited TCR-mediated T-cell proliferation with similar potency (IC₅₀ = 9-10 μM), and neither affected IL2-induced proliferation, underscoring the TCR specificity of both C10 forms (Supplementary Fig. 3a,b).

Collectively, these findings establish C10 as a highly selective inhibitor of TCR-mediated T-cell activation, with no detectable effects on B-cell activation or IL2-driven T-cell proliferation. In contrast, C6 and C8 exhibited broader inhibitory effects, making C10 the most promising lead compound for selective T-cell modulation.

### C8 and C10 inhibit TCR phosphorylation without affecting LCK kinase activity

To investigate the inhibitory effect of C8 and C10 on TCR signalling, we assessed their impact on TCR phosphorylation, a critical step in early T-cell signalling. Using JK T cells, we stimulated TCR signalling with an anti-hCD3ε antibody in the presence of DMSO (D), C6, C8, or C10. Anti-phospho-tyrosine immunoprecipitation followed by ζ chain immunoblotting revealed that C8 and C10 significantly reduced TCR phosphorylation by 50% compared to the DMSO control, while C6 had minimal or no impact (Fig. 4a). ZAP70 phosphorylation, a downstream target of LCK, was also reduced by 60% in the presence of C6, C8 and C10, further confirming inhibition of proximal TCR signalling (Fig. 4b). These findings suggest that C8 and C10 impair proximal TCR signalling by targeting an LCK-dependent mechanism.

**Fig. 4:**
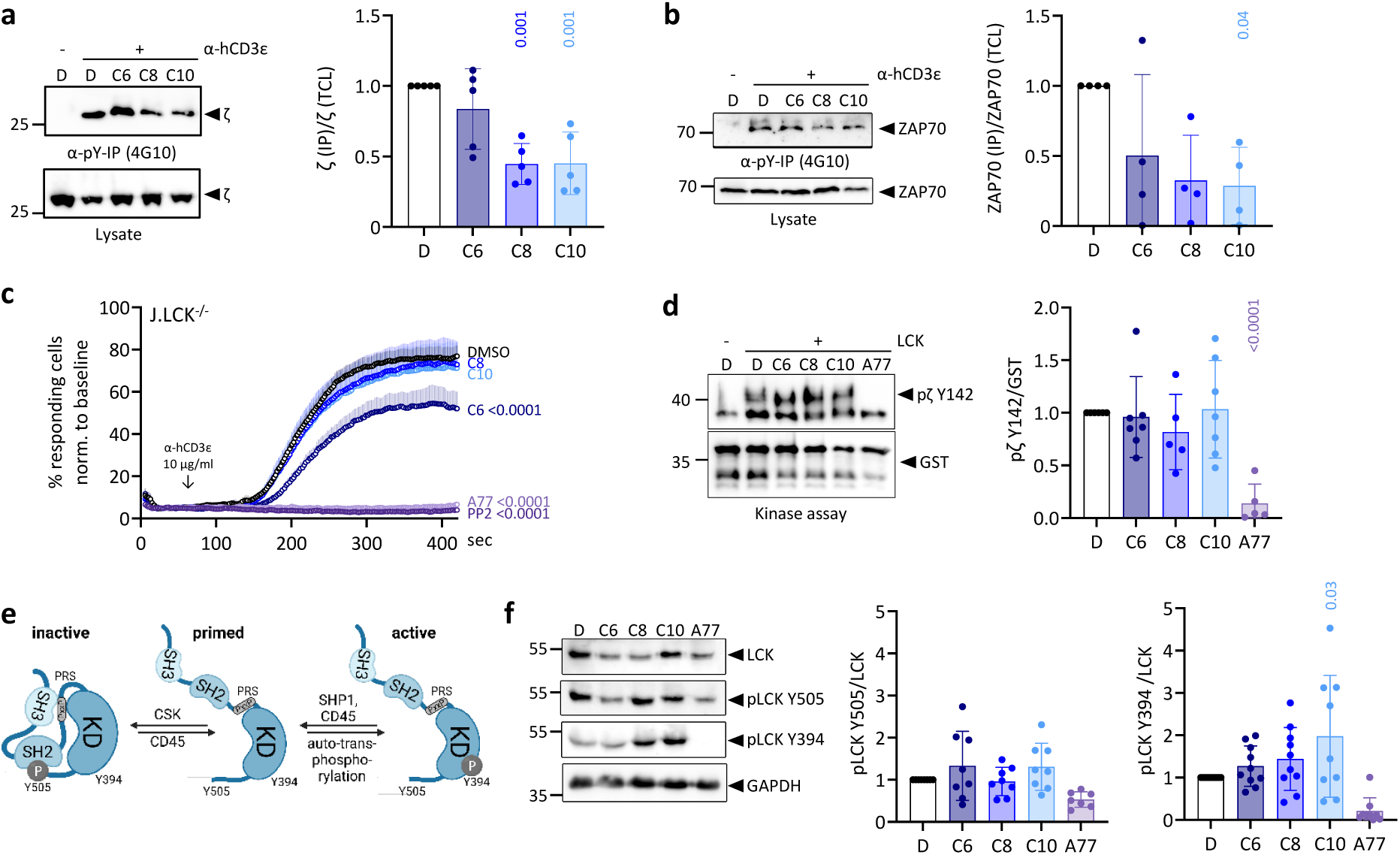
Novel compounds reduce early TCR signalling without affecting LCK activity. **a,b**, JK cells were incubated with 0.33% DMSO or 8 μM C6, C8 or C10 and stimulated with 5 μg/ml anti-hCD3ε for 5 min at 37 °C. Anti-phosphoY (4G10) immunoprecipitation (IP) was performed with directly coupled beads. Purified proteins and the lysate were loaded as control. Immunobloting was performed with the indicated antibodies. n = 4-5 independent experiments. **c**, J.LCK^-/-^ were stimulated with 10 μg/ml anti-hCD3ε, with baseline recorded for 60 s, in the presence of 0.33% DMSO or 8 μM C6, C8, C10, PP2, or A77. Percentage of Ca^2+^ fluxing cells over time is shown. n = 4-5 independent experiments. Means + SEM are shown. **d**, GST-coupled to the human cytoplasmic tail of ζ (GST-hcytCD3ζ) was incubated with ATP in the presence of 1% DMSO or 100 μM C6, C8 or C10, with or without recombinant full length human LCK for 15 min at 30 °C. **e**, Schematic representation of LCK regulation via conformational changes, phosphorylation, and intramolecular interactions. **f**, JK cells were treated with 0.33% DMSO or 8 μM C6, C8 or C10, and basal LCK phosphorylation at Y505 and Y394 was assessed by immunoblotting. Protein levels were quantified relative to the DMSO control and the GST or total lysate loading control. Each dot represents one independent experiment. Means ± SD are indicated. One-way ANOVA was used for statistical analysis.

To determine whether the reduction in phosphorylation was specifically due to LCK inhibition, we used JK LCK knockout (J.LCK^-/-^) cells, in which TCR signalling is highly compromised^43^. To optimize detection of Ca²⁺ influx in J.LCK^-/-^ cells, we first titrated the anti-hCD3ε antibody (Supplementary Fig. 4a). We then measured Ca²⁺ responses in J.LCK^-/-^ cells stimulated with 10 μg/mL anti-hCD3ε in the presence of DMSO, C6, C8, C10, PP2 (SFK inhibitor), or A77 (LCK inhibitor) (Fig. 4c). While all compounds did affect the Ca^2+^ response in JK WT cells (Supplementary Fig. 4b), C8 and C10 had no effect on Ca²⁺ signalling in J.LCK^-/-^ cells, confirming that their inhibition in T cells is LCK-dependent. In contrast, C6 still significantly reduced the percentage of responding cells by 60% compared to the DMSO control (Fig. 4c), supporting its unspecific mechanism of action. Interestingly, LCK inhibitor A77 fully blocked Ca^2+^ influx in the J.LCK^-/-^ cells, similarly to PP2, indicating off-target effects, most likely inhibiting the SFK FYN. These results confirm that C8 and C10 require LCK to exert their inhibitory effects, distinguishing them from broader SFK inhibitors.

To elucidate which domains of LCK are targeted by C8 and C10, we performed a kinase assay with human recombinant LCK in the presence of our compounds. None of the compounds affected the ability of LCK to phosphorylate the gluthathione S-transferase (GST)-fused ζ (Fig. 4d) or CD3ε cytoplasmic tails (Supplementary Fig. 4c), excluding that the compounds affect TCR phosphorylation by affecting the kinase function of LCK. The LCK kinase inhibitor A77 was included as positive control and completely blocked ζ and CD3ε phosphorylation by LCK (Supplementary Fig. 4c).

The SH3(LCK) domain is also involved in the regulation of the kinase’s activity through intramolecular conformational changes. In its inactive conformation, the regulatory tyrosine Y505 is phosphorylated by CSK, and binds the SH2 domain, while binding of the SH3 domain to the intramolecular proline-rich stretch (PRS) motif further stabilizes the closed-inactive LCK conformation^44^. De-phosphorylation of Y505 and phosphorylation of Y394 result in an open and active conformation of LCK (Fig. 4e)^45^. To assess whether the treatment with our compounds affects the intramolecular interaction of the SH3(LCK) domain with the PRS, we assessed the phosphorylation of Y505 and Y394 in basal conditions. Therefore, JK cells were treated with 8 μM of the compound for 1 h. Immunoblotting revealed an increase in Y394 phosphorylation after treatment with C10 (Fig. 4f), suggesting that it may disrupt SH3 domain-mediated stabilization of the inactive LCK conformation. However, the increase of pre-opened LCK under basal condition did not translate into increased basal TCR phosphorylation, which was unaffected by the compounds (Supplementary Fig. 4d). Furthermore, when stimulating the JK cells with an anti-hCD3ε antibody the phosphorylation of Y394 and Y505 remained unchanged in the presence of C8 or C10 (Supplementary Fig. 4e). The LCK kinase inhibitor A77 was used as a positive control and blocked the LCK kinase activity. These findings suggest that C10 may interfere with intramolecular interactions within LCK, favoring an open conformation (priming) without inducing constitutive activation.

Taken together, C8 and C10 impair early TCR signalling by reducing TCR and ZAP70 phosphorylation in an LCK-dependent manner, without inhibiting LCK kinase activity. C10 may also modulate LCK’s conformation, but this does not result in increased activity. These results highlight C10 as a selective and mechanistically distinct modulator of TCR signalling.

### C8 and C10 disrupt LCK recruitment to the TCR by targeting the SH3-CD3ε interaction

To determine whether C8 and C10 specifically disrupt the interaction between the SH3 domain of LCK and CD3ε, we performed pull-down (PD) assays and proximity ligation assays (PLA) to assess LCK recruitment to the TCR. We first conducted a PD assay using the unphosphorylated GST-tagged cytoplasmic domain of CD3ε (GST-cytCD3ε), which was bound to beads and incubated with JK cell lysates. SH3(LCK) binds the RK motif of CD3ε, while the adaptor protein NCK interacts via its SH3.1 domain with the PRS motif of CD3ε (Fig. 5a). In the presence of C8 and C10, LCK binding to CD3ε was reduced by 50% compared to the DMSO control, aligning with the 50% reduction in TCR phosphorylation observed in Fig. 4a (Fig. 5b). C6 had minimal impact, suggesting that its inhibitory effects on TCR signalling occur through a different mechanism. NCK binding to CD3ε remained unaffected by all compounds, confirming that C8 and C10 selectively disrupt the SH3(LCK)-RK interaction without broadly affecting SH3-mediated interactions, at least as assessed by this control (Fig. 5b).

**Fig. 5:**
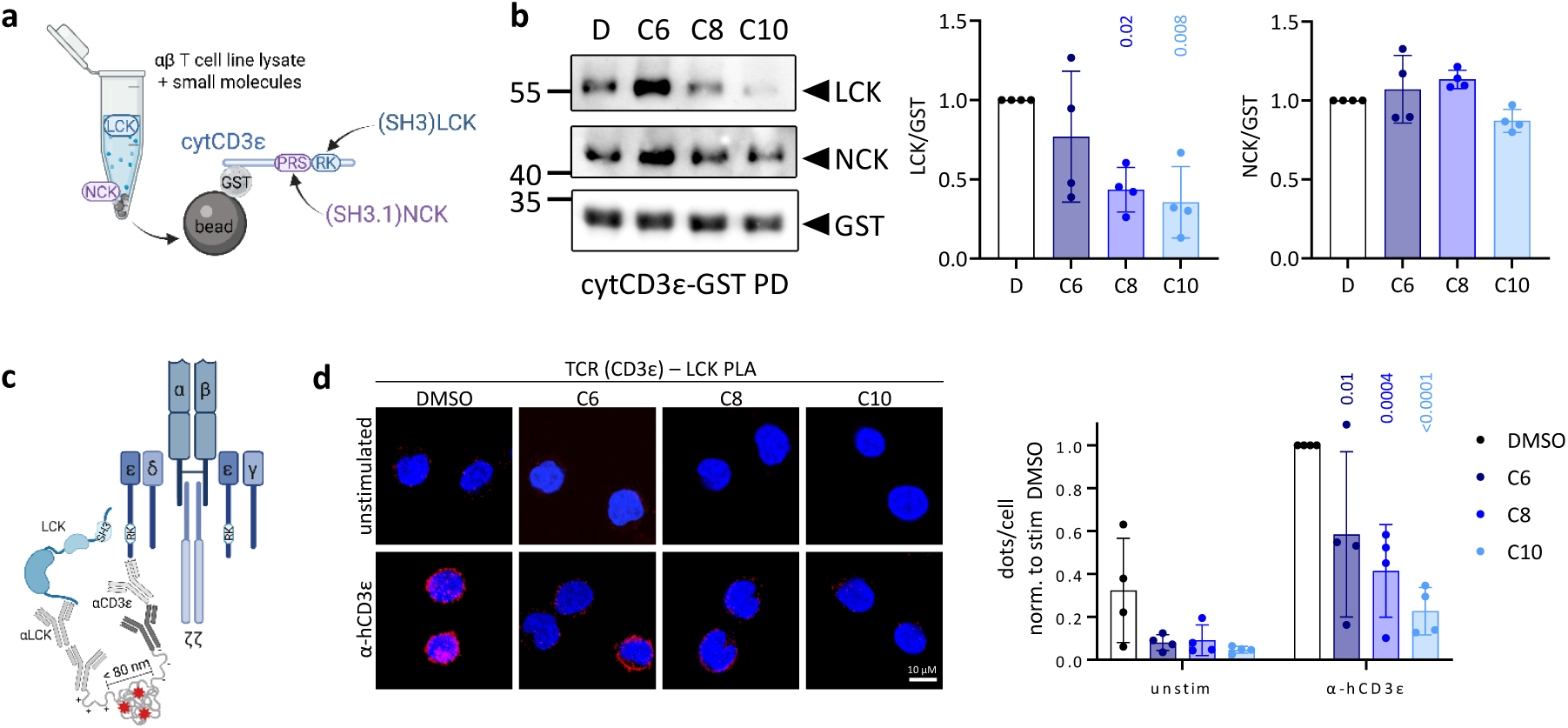
C8 and C10 impair LCK binding to CD3ε by disrupting SH3-mediated recruitment. **a**, Schematics of the assay performed in **b**. **b**, GST-cytCD3ε pull down assay in JK lysate in the presence of 1% DMSO or 100 μM C6, C8 or C10. Immunoblotting was performed using anti-LCK, -NCK and - GST antibodies. Band intensity was normalized to GST as a loading control and the DMSO control for each experiment. n = 4 independent experiments. Statistical analysis was performed using One-way ANOVA. **c**, Schematic of a TCR(CD3ɛ)-LCK proximity ligation assay (PLA). **d**, JK cells were either left unstimulated or stimulated with 5 µg/ml anti-hCD3ε for 5 min at 37 °C. PLA was performed between the TCR (CD3ε) and LCK. A red dot indicates a proximity closer than 80 nm. Images from one representative experiment are shown. Cell nuclei are stained with DAPI. Quantification of n= 4 independent PLA experiments. Dots per cell are normalized to the stimulated DMSO-treated sample for each experiment. Means ± SD are shown. Two-way ANOVA was used for statistical analysis.

To further validate this mechanism, we performed a PLA, which detects protein-protein proximities *in situ* within 80 nm by generating fluorescent signals in the presence of closely associated proteins (Fig. 5c). JK cells were left unstimulated or stimulated with anti-hCD3ε for 5 min in the presence of DMSO, C6, C8, or C10, and the proximity of CD3ε and LCK was assessed. Under basal conditions, all compounds slightly reduced TCR-LCK proximity, but the effect was not significant (Fig. 5d). Upon TCR stimulation, treatment with C6 reduced TCR-LCK proximity by 40%, treatment with C8 by 60% and treatment with C10 by 70%, compared to the DMSO control (Fig. 5d). These results confirm that C8 and C10 interfere with LCK recruitment to the TCR upon activation, supporting their role as targeted disruptors of the RK-LCK interaction.

Using biochemical and imaging-based approaches, we demonstrate that C8 and C10 selectively disrupt the SH3(LCK)-CD3ε interaction, providing strong evidence that they inhibit TCR signalling by interfering with LCK recruitment rather than by directly inhibiting its kinase activity.

### C10 selectively fine-tunes εCAR and TRuC T-cell activation

To further explore the therapeutic potential of our newly identified compounds, we assessed their ability to modulate CAR and TRuC T-cell activity. While CAR and TRuC T-cell therapies have demonstrated remarkable success in treating leukemia^46^, severe side effects such as CRS and neurotoxicity remain a significant challenge^13^. Therefore, selectively fine-tuning CAR and TRuC T-cell activation represents an attractive therapeutic strategy.

We have included five different CD19-targeting constructs in our panel. First the FDA-approved CAR construct, containing the 4-1BB-derived co-stimulatory domain and the ζ-derived signalling domain (BBζ). The signalling capacity of this construct does not dependent on the RK-LCK interaction. Then, a CAR containing CD3ε instead of ζ (BBε), and a CAR containing a CD3ε and ζ signalling domain (BBεζ) were used. The activity of these constructs is dependent on LCK recruitment via the RK motif^6,9^. Furthermore, we analyzed a BBεζ CAR containing the loss-of-function mutation RKAA within the RK motif (BBε^AA^ζ), and finally, we also included a TRuC construct^7^, where the single-chain variable fragment targeting CD19 is fused to CD3ε (Fig. 6a).

**Fig. 6:**
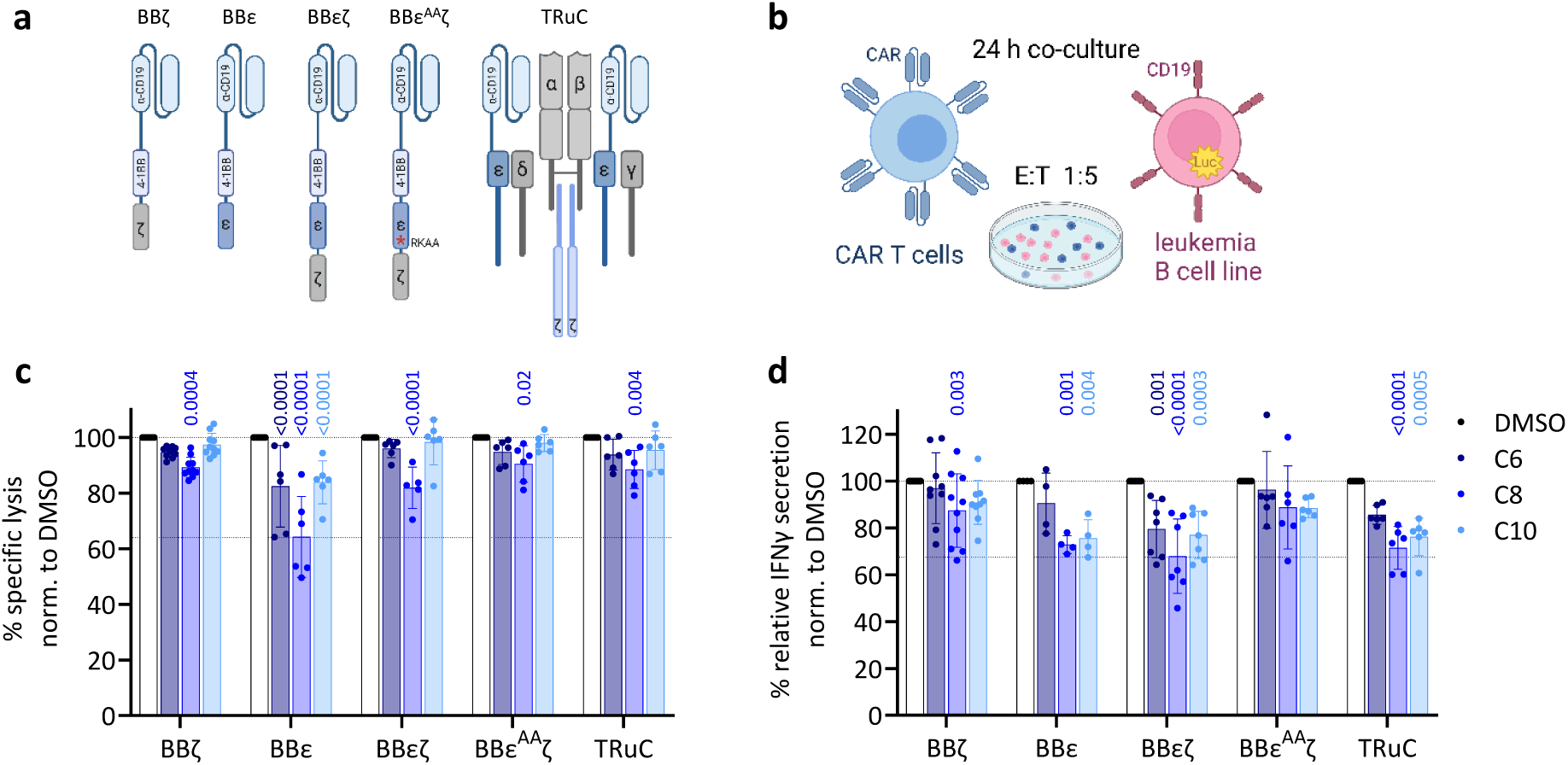
Selective modulation of cytotoxicity and cytokine secretion in CD3ε-containing CAR and TRuC T cells by SH3(LCK) inhibitors. **a**, Schematic representation of the anti-CD19 CARs and TRuC constructs used in this study. **b**, Primary human T cells, lentivirally transduced with the respective CAR or TRuC construct, were co-cultured with CD19^+^ luciferase-expressing Nalm6 cells in the presence of 0.33% DMSO or 8 μM of C6, C8 or C10 at an effector-to-target cell ratio of 1:5. **c**, Specific target cell killing was quantified by measuring the Nalm6 bioluminescence signal after 24 h of co-culture. **d**, Relative IFNγ secretion was assessed by ELISA after 24 h of co-culture. Each dot represents one independent experiment. Data points are normalized to the DMSO control for each CAR/TRuC construct to highlight compound-specific effects. n = 2-10 HDs in 1-2 independent experiments each. Means ± SD are shown. One-way ANOVA was used for statistical analysis.

We lentivirally transduced primary human T cells with vectors encoding these different constructs, and transduction efficacy was assessed by flow cytometry (Supplementary Fig. 5a-d). To assess CAR/TRuC functionality, T cells were co-cultured with CD19-expressing Nalm6 leukemia cells (Fig. 6b). After 24 h, tumour cell killing was evaluated by bioluminescence-based cytotoxicity assays, while IFNγ secretion was assessed by ELISA. Since different CAR and TRuC constructs exhibit varying cytotoxicity and cytokine secretion upon stimulation with CD19^+^ target cells (Supplementary Fig. 5e), we normalized all data to the DMSO control for each construct to highlight the specific effects of our compounds on each construct. All three compounds reduced cytotoxicity of the BBε CAR (LCK-dependent), indicating that disrupting LCK recruitment impairs killing efficiency. The killing capacity of the BBεζ CAR was only reduced by C8, suggesting that the presence of ζ compensates for the inhibitory effect of the compounds. In line, the killing capacity of the TRuC was hardly affected, also suggesting some compensation by the additional CD3 chains (Fig. 6c). As expected, the BBζ and BBεAAζ CARs (LCK-independent controls) were unaffected by any compound, confirming that the observed effects are specific to LCK-mediated signalling.

To rule out off-target effects on target cells, we performed luciferase-based proliferation assays with Nalm6 and other leukemia cell lines. While C8 inhibited leukemia cell proliferation, C10 had no direct effect, reinforcing its T-cell specificity (Supplementary Fig. 5f,g). We next evaluated IFNγ secretion. C8 and C10 significantly reduced IFNγ secretion of the BBε CAR, the BBεζ CAR, and the TRuC, while hardly affecting the BBζ and BBε^AA^ζ control constructs (Fig. 6d).

These results demonstrate that C10 selectively modulates CD3ε-containing CAR and TRuC T-cell activation, reducing IFNγ secretion in an LCK-dependent manner. This highlights the therapeutic potential of this innovative of T-cell modulator to control undesired CAR/TRuC T-cell cytokine secretion.

### C10 promotes a central-memory-like phenotype in εCAR T cells

Long-term persistence of CAR T cells is critical for durable therapeutic responses, with central memory (TCM)-like subsets displaying enhanced survival and anti-tumour efficacy *in vivo*^47^. Given the dependence of CD3ε-containing CARs on LCK-mediated signalling, we investigated whether C10 treatment could influence CAR T-cell differentiation by promoting a TCM-like phenotype while limiting terminal effector subsets. To assess this, BBε CAR T cells were cultured in the absence of any compound (x), in the presence of DMSO (D), or 2, 4, or 8 μM C10 for 13 days, starting two days post-transduction (Fig. 7a). Transduction efficiency and CAR expression, assessed on day 7, were comparable across all treatment conditions (Supplementary Fig. 6a-d), and the CD4/CD8 ratio remained unchanged (Supplementary Fig. 6e). While low-doses of C10 did not affect the expansion potential of BBε CAR T cells, the highest dose reduced expansion by approximately half relative to control conditions, although the effect was not statistically significant among donors (Fig. 7b). Phenotypic analysis via flow cytometry revealed a dose-dependent increase in T_CM_-like CAR T cells and a concurrent reduction in effector memory (T_EM_) and terminally differentiated effector (T_EMRA_) subsets at basal conditions (Fig. 7c). Upon 24-hour co-culture with CD19+ Nalm6 cells, these effects were further accentuated, with 8 μM C10 yielding a CAR T-cell phenotype closely resembling mock-transduced T cells. Importantly, BBζ CAR T cells, which lack CD3ε signalling, were unaffected by C10, confirming the compound’s specificity for CD3ε-mediated signalling (Supplementary Fig. 6f).

**Fig. 7:**
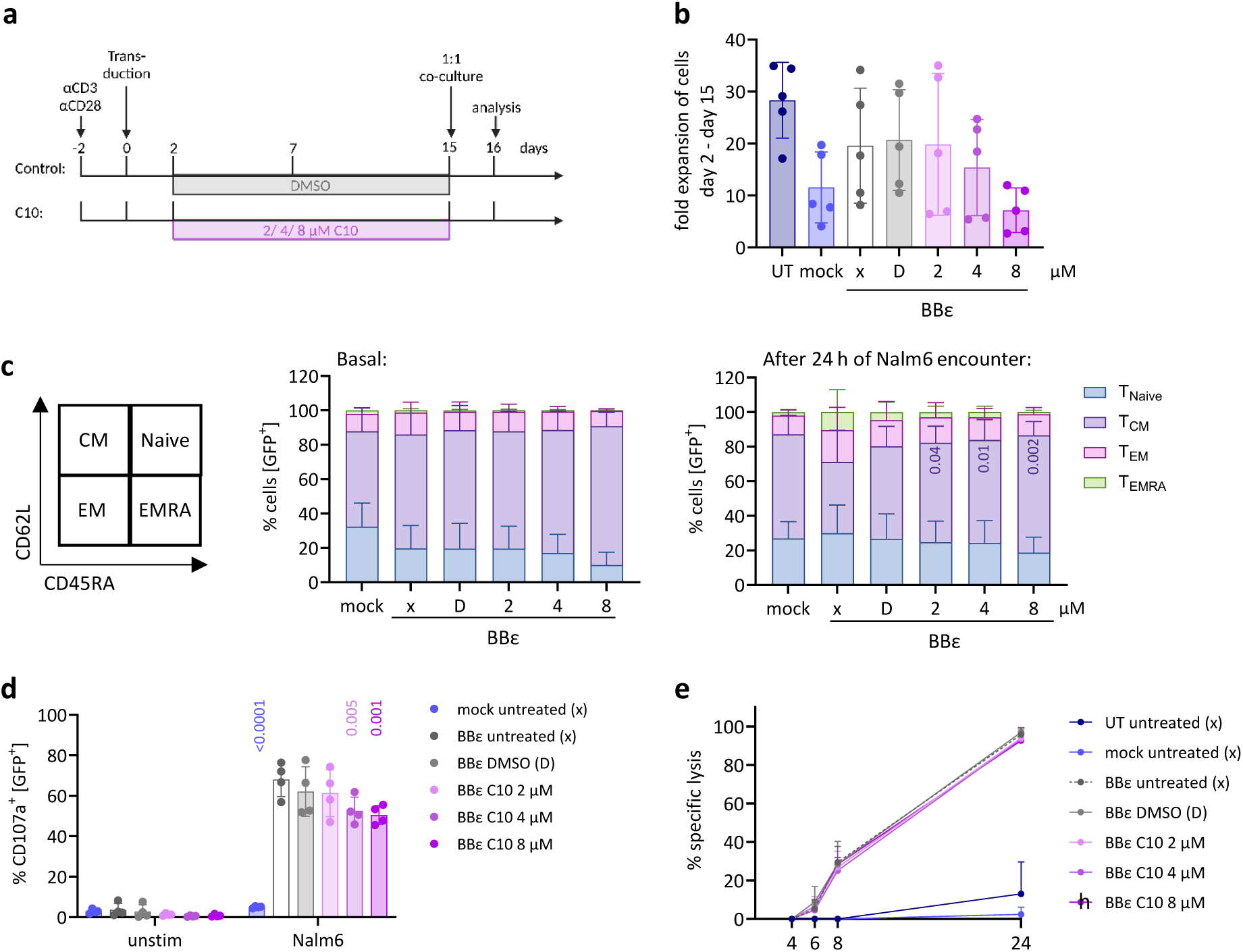
Transient C10 treatment promotes a central memory–like phenotype in BBε CAR T cells without impairing cytotoxicity. **a**, Timeline of CAR-T-cell manufacturing, including treatment with 2, 4 or 8 μM of C10 for 13 days. DMSO-treatment was used as control. **b**, Fold expansion of cells between day 2 and day 15 of the experiment. n = 5 HDs. **c**, Phenotypic analysis of BBε CAR-T cells by flow cytometry, using anti-hCD62L and anti-hCD45RA antibodies. The percentage of respective populations is shown under basal conditions and after 24 h of co-culture with Nalm6 target cells. n = 5-9 HDs. **d**, Degranulation of CAR-T cells was assessed after 3 h of stimulation with Nalm6 target cells by flow cytometry using an anti-hCD107a antibody. n = 4 HDs **e**, Specific target cell killing was measured by assessing the Nalm6 bioluminescence signal after 4, 6, 8 and 24 h of co-culture with CAR-T cells. n = 4 HDs. Each dot represents one HD. Means ± SD are shown. Statistical analysis was performed using one-way (b, c) or two-way (d) ANOVA comparing to untreated BBε CAR-T cells.

To ensure that C10 pretreatment did not compromise CAR T-cell function, we assessed activation, cytokine production, and cytotoxicity following a 24-hour co-culture with Nalm6 cells. C10-pretreated cells upregulated CD69, CD25, and 4-1BB to levels comparable to controls (Supplementary Fig. 6g), and secreted IL2, IFNγ, and TNF effectively (Supplementary Fig. 6h). While a slight reduction in CD107a degranulation was observed at higher C10 concentrations (Fig. 7d), cytotoxicity measured by a luciferase-based killing assay remained intact, with complete target cell clearance across all conditions (Fig. 7e). Together, these findings position C10 as a novel pharmacological tool to reprogram εCAR T-cell differentiation toward a T_CM_-like state without compromising effector function, offering a promising strategy to enhance CAR T-cell manufacturing and clinical performance.

## Discussion

Fine-tuning T-cell activation is crucial for immunotherapy, particularly in the treatment of autoimmune diseases, GVHD and CAR T-cell therapy. A central step of T-cell activation is the recruitment of LCK to the TCR, mediated by the non-canonical interaction between its SH3 domain and the RK motif of CD3ε^6^. Our study reinforces the functional relevance of this interaction not only in native TCR signalling, but also in engineered constructs such as CAR and TRuC T cells, where CD3ε signalling has emerged as a powerful component^6,30^. This underexplored mechanism provides a compelling target for pharmacologic intervention.

Current strategies to modulate T-cell activation primarily rely on broad kinase inhibitors, which often cause systemic immunosuppression and off-target effects. While LCK is a key target for drug development, existing inhibitors exclusively focus on its highly conserved kinase domain, leading to unintended inhibition of other SFKs such as FYN, LYN, SRC and YES^4,5^. To address this limitation, we introduce C10 as a first-in-class small-molecule modulator that selectively disrupts the SH3(LCK)-RK interaction and thereby impairs TCR signalling at the level of LCK recruitment, without inhibiting its catalytic activity.

C10 dampened TCR-mediated T-cell activation, proliferation, and cytokine secretion, while sparing IL2-driven proliferation and B-cell responses, demonstrating high specificity for the TCR signalling axis. Mechanistically, it reduced TCR and ZAP70 phosphorylation without affecting the catalytic activity of LCK, distinguishing it from classical kinase inhibitors. Pull-down and proximity ligation assays confirmed selective disruption of LCK recruitment to CD3ε. By targeting a protein-protein interaction rather than enzymatic function, C10 enables precise and selective modulation of T-cell activation, offering a promising alternative to conventional LCK inhibition.

The therapeutic potential of targeting protein-protein interactions in early TCR signalling has been previously explored. One study reported the development of AX-024, a small-molecule inhibitor that binds the SH3.1 domain of the adaptor protein NCK, disrupting its interaction with the PRS motif of CD3ε and thereby reducing TCR signalling^48^. AX-024 demonstrated protective effects in mouse models of psoriasis, allergic asthma and multiple sclerosis, highlighting the feasibility of this approach. Given that RK motif mutations have a more pronounced impact on T-cell activation than PRS motif mutations^6^, SH3(LCK)-targeting inhibitors such as C10 may even offer superior therapeutic benefits.

Importantly, reducing LCK recruitment to the TCR does not fully abolish signalling but instead dampens it^6^, making our SH3(LCK)-targeting compounds immunomodulatory rather than immunosuppressive. By reducing, rather than eliminating, TCR signalling, C10 may offer precise control over T-cell activity, which is particularly relevant in clinical settings such as GVHD. GVHD can occur following allogeneic hematopoietic stem cell transplantation in leukaemia patients, where allogeneic donor T cells are activated by host APCs, triggering life-threatening inflammation and severe tissue damage^49,50^. Acute GVHD is a T cell-dependent pathology, and therapies aim to reduce T-cell activity to control its severity while still allowing T cells to eliminate leukemic cells, thus preserving the beneficial graft-versus-leukaemia (GVL) effect^51^. Our *in vitro* results demonstrate that SH3(LCK)-targeting compounds significantly reduce CD69 upregulation and T-cell proliferation in allogeneic stimulation assays, supporting their potential as potential GVHD therapeutics. Additionally, C10 could then be explored in disease models like diabetes type-1, asthma, psoriasis, rheumatoid arthritis, transplant rejections or multiple sclerosis.

Beyond immune modulation, our findings highlight the potential of SH3(LCK)-targeting compounds in CAR T-cell therapy. Excessive CAR signalling is a major challenge, often leading to CRS and premature T-cell exhaustion^11,52–55^. While CAR and TRuC T-cell therapies have demonstrated remarkable success in treating hematologic malignancies, continuous efforts are underway to refine these approaches and improve safety and efficacy. Incorporating the CD3ε chain as a signalling domain in CAR constructs enhances anti-tumour efficacy both *in vitro* and *in vivo*^6,8–11^.

The selective inhibition of LCK recruitment by C10 provides a pharmacological tool to fine-tune the activation of CD3ε-containing CARs and TRuCs, reducing both cytotoxicity and IFNγ secretion. These effects were not observed in ζ-based CAR constructs, reinforcing the specificity of C10 for CD3ε-mediated signalling. Given that excessive T-cell activation can lead to severe side effects such as CRS and neurotoxicity, a pharmacological approach to transiently dampen CAR and TRuC activity may offer a safer and more controlled alternative to genetic modifications that permanently alter signalling strength. Beyond modulating acute T-cell activation, C10 also influenced the differentiation of CAR T cells, promoting a central-memory-like phenotype while reducing effector memory and terminal effector subsets. T_CM-_enriched CAR T-cell populations have been associated with enhanced persistence, reduced exhaustion, and superior long-term therapeutic efficacy^47^. Importantly, C10-pretreated CAR T cells retained full activation potential and cytotoxic function, indicating that the compound modulates differentiation without compromising effector capacity. This suggests that C10 could be incorporated into CAR T-cell manufacturing protocols to generate long-lived, therapeutically potent T cells with improved persistence *in vivo*. Previous studies have already shown that tyrosine kinase inhibitors, such as dasatinib, can serve as pharmacologic tools to modulate CAR T-cell activity. Specifically, they have been shown to reduce early exhaustion and preserve stem-like properties of CAR T cells, and diminish CRS severity in mouse models^56–59^. Our data suggest that C10 could be explored in a similar capacity, providing an additional layer of control over CAR T-cell activity.

While these findings establish C10 as a promising candidate for selective T-cell modulation, further structural optimization is recommended to improve its pharmacokinetics and solubility before investigating *in vivo* efficacy.

In conclusion, this study identifies a first-in-class T-cell modulator targeting the SH3 domain-mediated recruitment of LCK to the TCR, providing a precise and reversible mechanism to regulate T-cell activation. Our findings highlight the potential of this innovative T-cell modulator not only for the treatment of a diverse range of autoimmune and inflammatory diseases, but also as a pharmacologic tool to improve the efficacy and safety of CAR T-cell therapy.

## Acknowledgements

We thank Marina Freudenberg for performing an initial test experiment and Kerstin Fehrenbach for excellent technical support. M.K., W.W.S, N.K. and S.M. are supported by the German Research Foundation (DFG) through BIOSS - EXC294 and CIBSS - EXC 2189. S.M. and N.K. are supported by DFG via SFB1479 (Project ID: 441891347 - P15 to S.M., P03 to N.K.), SFB1160 (Project ID: 256073931 - B01 to S.M, B09 to N.K.) and MI1942/5-1 (Project ID: 501436442 to S.M.). W.S. is supported by DFG via SFB1381 (Project ID: 403222702 - A9). This project was funded by the German Cancer Aid (70116490 to N.K.).

## Author contributions

N.M.W., T.R., L.W., M.Z., A.M.S., and S.H. performed experiments and data analysis. P.C. and P.L.M. performed the docking-based virtual screening and *in silico* characterization. N.M.W., P.C., T.R., A.M.S., A.K., M.K., M.C., F.A.H., W.W.S., N.K., P.L.M., and S.M. provided intellectual input. S.M. and P.L.M. conceived the project and designed the study. N.M.W. and S.M. wrote the manuscript with the input of all authors.

