## Supplemental File for "Controlling TCR and CAR activation by targeting LCK recruitment with a first-in-class small-molecule inhibitor"

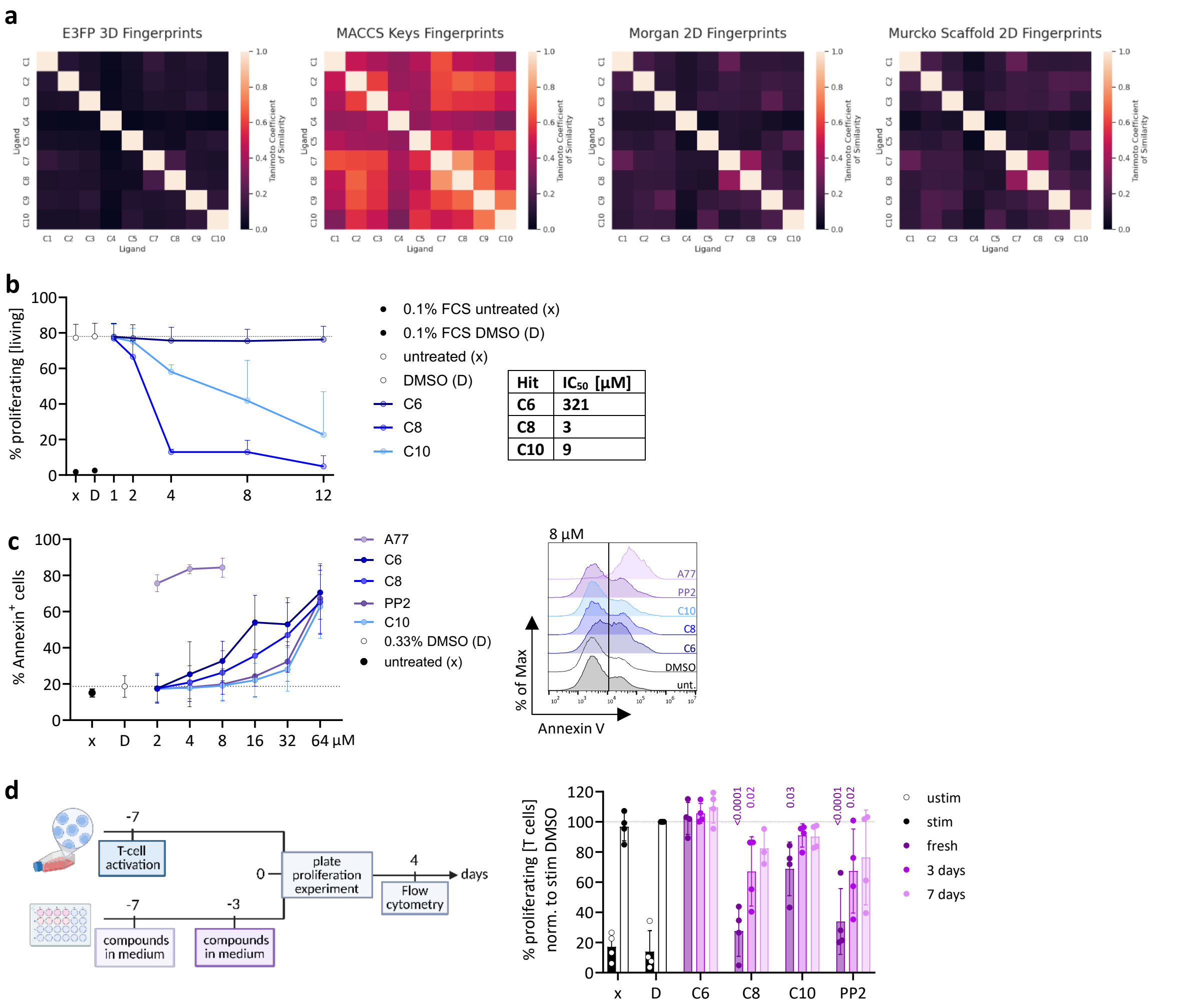


**Supplementary Fig. 1: Fingerprint similarity matrix of screening compounds and evaluation of compound toxicity and stability at physiological temperature. a,** Tanimoto coefficiancy of similarity heatmaps for all ten SH3(LCK)-targeting candidate compounds (C1–C10) calculated using four molecular fingerprinting methods: E3FP 3D fingerprints, MACCS keys, Morgan 2D fingerprints, and Murcko scaffold fingerprints. **b,** JK cells were incubated in 0.1% FCS medium (negative control) or in 10% FCS medium and either left untreated (x), treated with 0.33% DMSO (D) or with different concentrations of C6, C8 or C10. After 72 h, the percentage of proliferating cells was assessed by flow cytometry. Means + SD are shown. n = 3 independent experiments. The $IC_{50}$ of each compound for inhibiting proliferation is shown in the table. **c,** Primary human T cells were either left untreated (x) or treated with 0.33% DMSO or different concentrations of C6, C8, C10, PP2, and A77. Annexin V staining was used to assess cell viability after 24 h by flow cytometry (left). n = 3-8 healthy donors (HDs). Means ± SD are shown. The dotted line represents baseline of DMSO-induced cell death. Representative histograms from one HD are shown (right). **d,** Primary human T cells were either left unstimulated or stimulated with 5 µg/ml anti-hCD3ε for 96 h. Cells were left untreated (x), treated with 0.33% DMSO, 8 µM of fresh C6, C8, C10 or PP2, or with medium containing 8 µM of C6, C8, C10, or PP2 that had been kept for three or seven days at 37 °C. The percentage of proliferating $CD8^+$ or $CD4^+$ T cells was assessed by flow cytometry. Data are normalized to the stimulated DMSO control. Means ± SD are shown. Each dot represents one HD; n = 4 HDs. Two-way ANOVA was used for statistical analysis.

**a**

**Human SH3(LCK)**: LQDNLVIALHSYEPSHDGDLGFEKGEQLRILEQSGEWWKAQSLTTGQEGFIPFNFVAKANS

**Murine SH3(LCK)**: LQDNLVIALHSYEPSHDGDLGFEKGEQLRILEQSGEWWKAQSLTTGQEGFIPFNFVAKANS

**b**

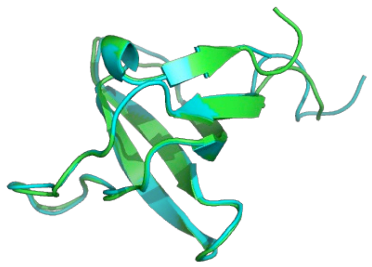

**Supplementary Fig. 2: Structural alignment of murine and human SH3(LCK) domains. a,** Amino Acid sequence of human and murine SH3(LCK). **b,** Ribbon overlay of the murine SH3(LCK) AlphaFold-predicted model (cyan) and the experimentally resolved human SH3(LCK) structure (green).

**a**

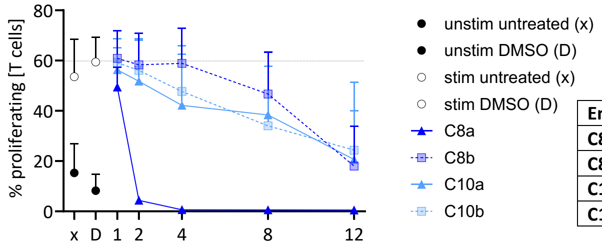


| Enantiomer | $IC_{50}$ [µM] |
|---|---|
| **C8a** | 1 |
| **C8b** | 17 |
| **C10a** | 9 |
| **C10b** | 10 |

**b**

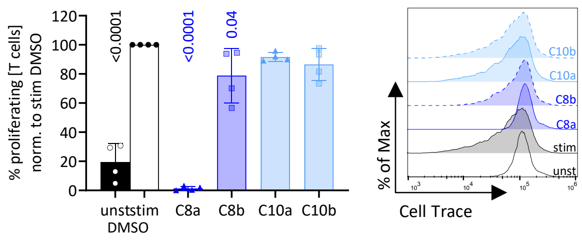


**Supplementary Fig. 3: Impact of C8 and C10 enantiomers on human T-cell proliferation. a,** Proliferation of $CD8^+$ and $CD4^+$ primary human T cells was assessed by flow cytometry after stimulation with 5 µg/ml anti-hCD3ε in the presence of 0.33% DMSO or the C8 or C10 enantiomers for 96 h. n = 4 HDs. Means + SD are shown. $IC_{50}$ values for the enantiomers calculated based on the proliferation data. **b,** Primary human T cells were left unstimulated or stimulated with 100 ng/ml IL2 for 96 h, and proliferation was assessed by flow cytometry. n = 4 HDs. Means ± SD are shown. One-way ANOVA was used for statistical analysis.

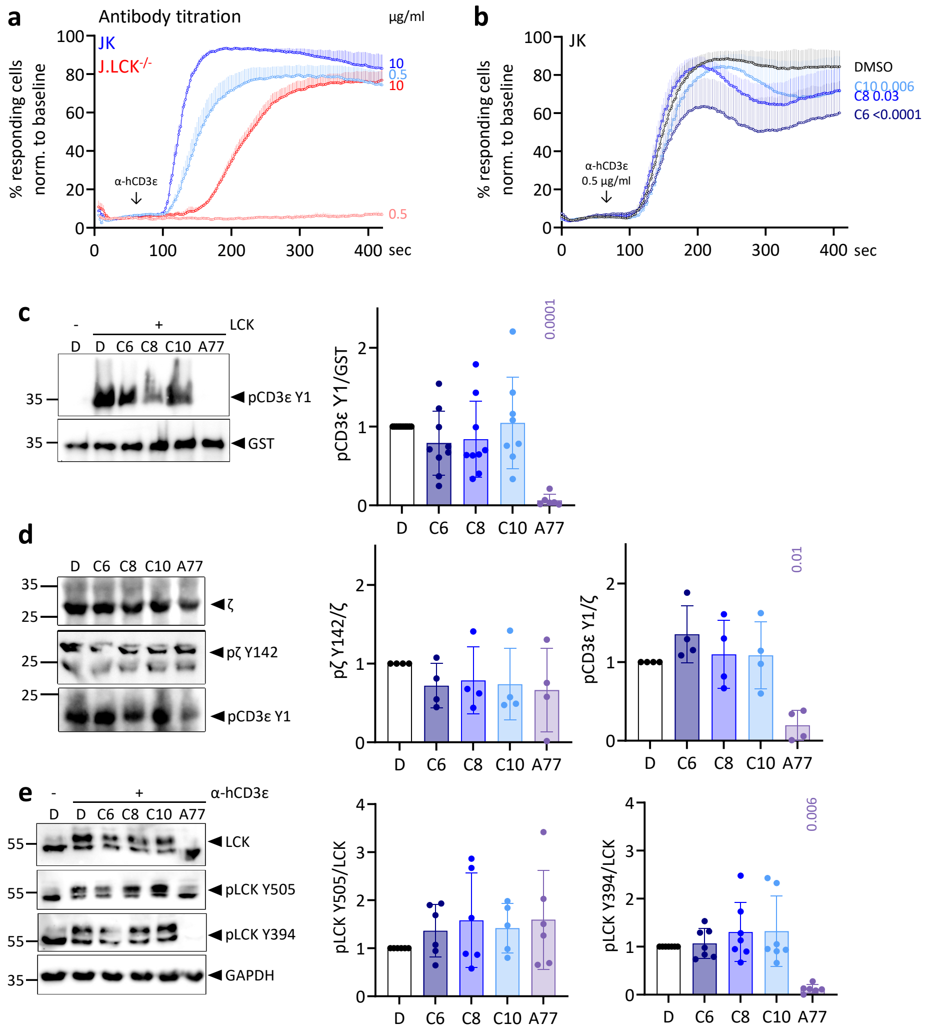


**Supplementary Fig. 4: Modulation of early TCR signaling and phosphorylation events by C6, C8, and C10 in JK cells. a,** JK and JK after CRISPR/Cas9 KO of LCK (J.LCK$^{-/-}$) were stimulated with 10 or 0.5 µg/ml anti-hCD3ε antibody after recording baseline for 60 sec. n = 2-5 independent experiments. **b**, JK WT cells were stimulated with 0.5 µg/ml anti-hCD3ε, after baseline recording for 60 sec, in the presence of 0.33% DMSO or 8 µM C6, C8 or C10. n = 3 independent experiments. Means +SEM are shown. **c**, hcytCD3ε-GST was incubated with ATP in the presence of 1% DMSO or 100 µM C6, C8 or C10 with or without recombinant full length human LCK for 15 min at 30 °C. **d**, JK cells were treated with 0.33% DMSO or 8 µM C6, C8 or C10. Basal Zeta (Y142) and CD3ε (Y1) phosphorylation was assessed by immunoblotting. **e**, JK cells were incubated with 0.33% DMSO or 8 µM C6, C8 or C10 and left unstimulated or stimulated with 5 µg/ml anti-hCD3ε for 5 in at 37 °C. Protein levels were quantified relative to the DMSO control and normalized to the GST or total lysate loading control. Each dot represents one independent experiment. Means ± SD are shown. One-way ANOVA was used for statistical analysis.

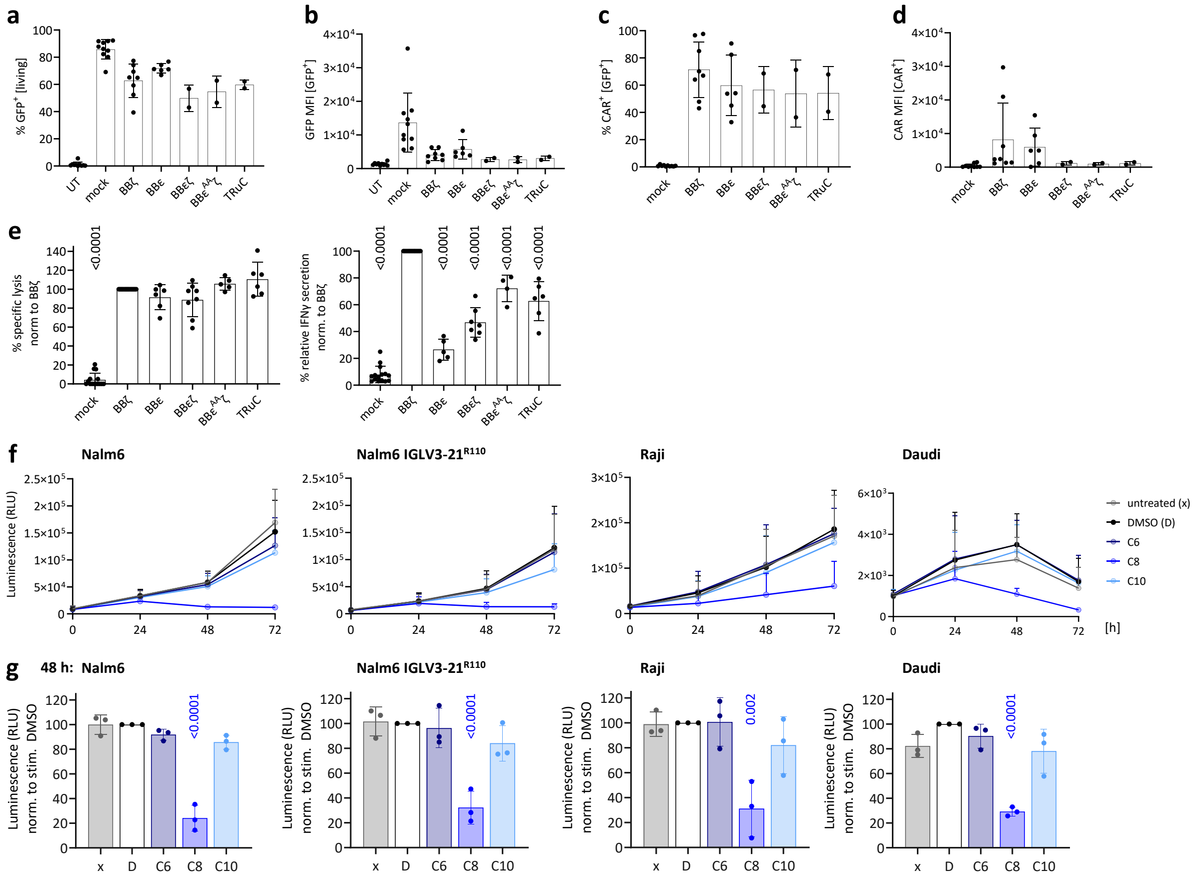


**Supplementary Fig. 5: Cytotoxicity and relative IFNγ secretion of different CAR and TRuC T cells and proliferation of leukemic B-cell lines.** Flow cytometry analysis of primary human T cells lentivirally transduced with the indicated constructs. Untransduced T cells (UT) were used as negative control. **a**, Percentage of GFP$^+$ cells. **b**, MFI of GFP in GFP$^+$ cells, respectively. **c**, Percentage of CAR$^+$ cells. **d**, MFI of CAR in CAR$^+$ cells. Each dot represents one HD. n = 2-10 HDs. **e**, CAR and TRuC T-cells were co-cultured with CD19$^+$, luciferase expressing Nalm6 cells with an effector-to-target cell ratio of 1:5. Specific target cell killing was assessed by measuring the Nalm6 bioluminescence after 24 h. Relative IFNγ secretion was measured by ELISA after 24 h of Nalm6 co-culture. Each dot represents one independent experiment. Data points are normalized to the BBζ CAR-T cells in each experiment. n = 4-6 HDs in 1-2 independent experiments each. **f**, Proliferation of luciferase-expressing leukemic B-cell lines was assessed by measuring bioluminescence over time in the presence of 0.33% DMSO or 8 μM of C6, C8 or C10. Nalm6 IGLV3-21$^{R110}$ express a distinctoncogenic BCR. **g**, The 48 h datapoints from **f** was normalized to the DMSO control. Each dot represents an independent experiment. n = 3. Means ± SD are shown. One-way ANOVA was used for statistical analysis.

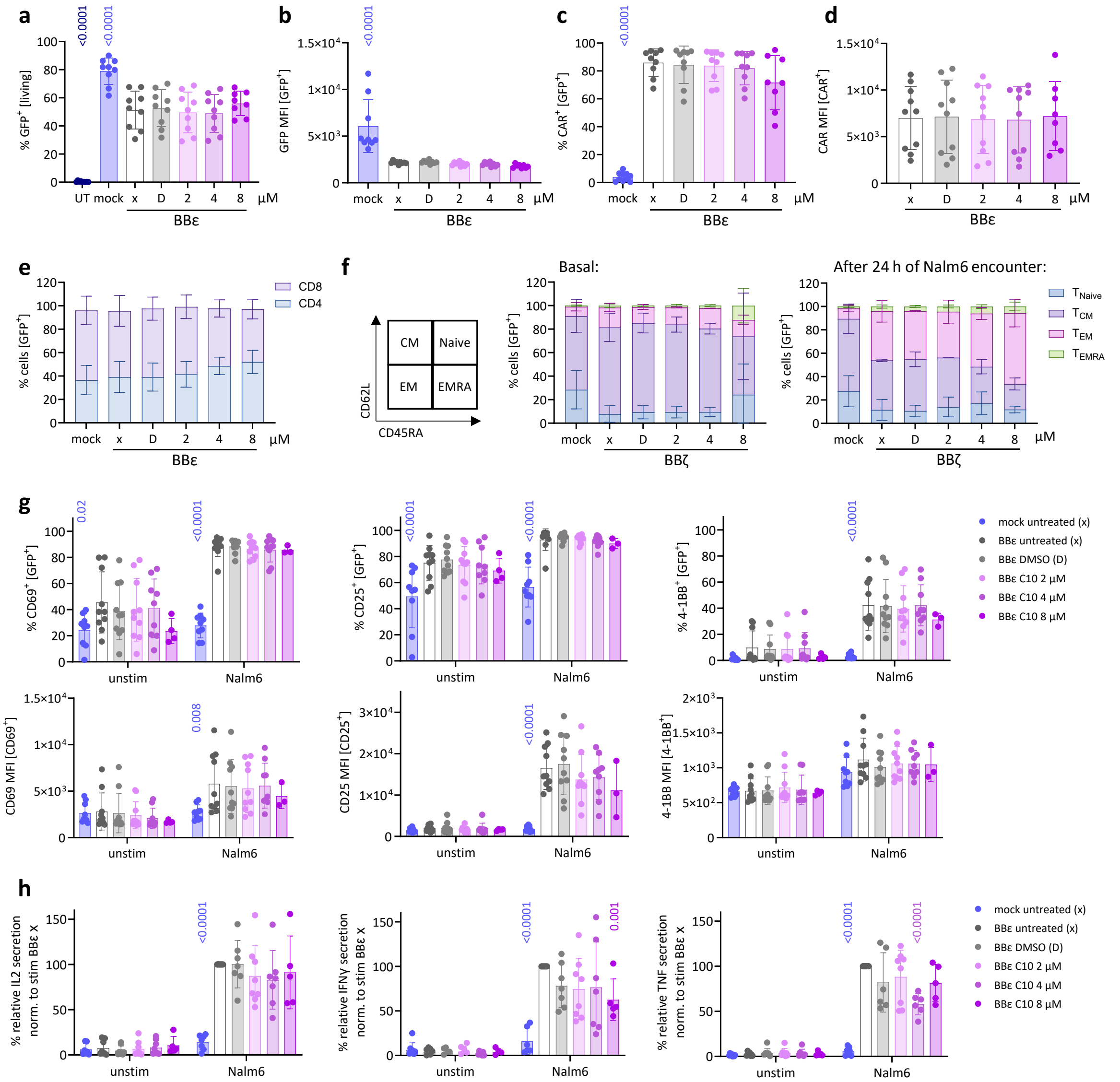


**Supplementary Fig. 6: Characterization of rested CAR-T Cells: transduction efficiency, differentiation, and activation potential. a**, Percentage of GFP⁺ cells. **b**, MFI of GFP in GFP⁺ cells. **c**, Percentage of CAR⁺ cells. **d**, MFI of CAR in CAR⁺ cells. **e,** CD4/CD8 ratio of GFP⁺ cells was assessed by flow cytometry. n = 3-6 HDs. **f**, Differentiation analysis of BBζ CAR-T cells by flow cytometry using anti-hCD62L and anti-hCD45RA antibodies. The percentage of respective populations is shown under basal conditions and after 24 h of co-culture with Nalm6 target cells. n = 2 HDs. **g**, Activation marker upregulation was assessed by flow cytometry using anti-hCD69, anti-hCD25, and anti-h41BB antibodies under basal conditions and after 24 h of co-culture with Nalm6 target cells. n = 3-10 HDs. **h**, The supernatants after 24 h of co-culture with Nalm6 cells were assessed for cytokines IL2, IFNγ and TNF via ELISA. All data were normalized to the Nalm6-stimulated, untreated BBε CAR-T cells. n = 5-8 HDs. Each dot represents one HD. Means ± SD are indicated. Statistical analysis was performed using one-way (a-f) or two-way (g, h) ANOVA comparing to untreated BBε/BBζ CAR-T cells.
